# Homeostatic maintenance and age-related functional decline in the *Drosophila* ear

**DOI:** 10.1101/764670

**Authors:** Alyona Keder, Camille Tardieu, Liza Malong, Anastasia Filia, Assel Kashkenbayeva, Jonathan E. Gale, Mike Lovett, Andrew P. Jarman, Joerg T. Albert

**Affiliations:** Ear Institute, University College London, 332 Gray’s Inn Road, London WC1X 8EE, UK; National Heart and Lung Institute, Imperial College London, Guy Scadding Building, Dovehouse 9 Street, London, SW3 6LY, UK; Centre for Discovery Brain Sciences, Edinburgh Medical School, University of Edinburgh, Edinburgh EH8 9XD, Scotland, UK; Centre for Mathematics and Physics in the Life Sciences and Experimental Biology (CoMPLEX), University College London, Gower Street, London WC1E 6BT, UK; The Francis Crick Institute, 1 Midland Road, London NW1 1AT, UK; Department of Cell and Developmental Biology, University College London, Gower Street, London WC1E 6DE, UK

## Abstract

The widespread loss of hearing is one of the major threats to future wellbeing in ageing human societies. Amongst its various forms, age-related hearing loss (ARHL) carries the vast bulk of the global disease burden. The causes for the terminal decline of auditory function, however, are as unknown as the mechanisms that maintain sensitive hearing before its breakdown. We here present an in-depth analysis of maintenance and ageing in the auditory system of the fruit fly *Drosophila melanogaster*. We show that *Drosophila*, just like humans, display ARHL and that their auditory life span is homeostatically supported by a set of evolutionarily conserved transcription factors. The transcription factors Onecut (closest human orthologues: ONECUT2, ONECUT3), Optix (SIX3, SIX6), Worniu (SNAI2) and Amos (ATOH1, ATOH7, NEUROD1) emerged as key regulators acting upstream of core sensory genes, including components of the fly’s molecular machinery for auditory transduction and amplification.

## Introduction

A surface calm can be misleading. All living things, from unicellular amoeba to neurons in the human brain, require continual maintenance and the constant flow of their seemingly equable physiological operations is in fact the product of complex homeostatic networks. All life, it has been said, needs to run to stand still. As with many things, the underlying homeostasis machinery remains mostly unappreciated until it breaks down. A most pertinent example of such a breakdown are the hearing impairments that affect about 1.23 billion people worldwide, corresponding to one sixth of the world’s total population ^1^. The aetiology of hearing loss is diverse, including infections, ototoxic drugs, noise trauma, as well as genetic factors. The arguably single most important cause, however, is age. Age-related hearing loss (ARHL) carries the vast bulk of the global disease burden, but its scientific understanding is poor and no treatments, neither preventive nor curative, are currently in sight ^2^.

Over the past few decades, gene discovery studies using mouse models have identified numerous candidate genes for human deafness ^3–9^. Three recent larger scale screens have brought the total number of candidate hearing loss genes for humans to 111 ^10–12^. Yet, the underlying mechanisms and molecular networks of ARHL, and particularly the maintenance of hearing throughout the lifespan, have remained elusive. We here use the auditory system of the fruit fly to shed some light on these issues.

The auditory systems of vertebrates and *Drosophila* share marked similarities; these include (i) the fundamental mechanisms of auditory transduction ^13^ and amplification ^14,15^, (ii) the general architecture of neuronal pathways from the ear to higher-order centres in the brain ^16^ and, finally, (iii) conserved families of proneural genes that control hearing organ development, such as e.g. *ato* in flies and Atoh1 in mice (or ATOH1 in humans) ^17,18^. The high degree of similarity, and partially near identity, between the ears of *Drosophila* and vertebrates (including mammals) has recommended the fly as a powerful model to study fundamental aspects of human hearing and deafness ^19^.

Many *Drosophila* hearing genes have been identified ^19–21^, but so far no study has explored the flies’ hearing across their life course. We found that the ears of fruit flies also display ARHL. Going one step further we set out to identify those homeostatic regulators that maintain the fly’s sensitive hearing before the onset of ARHL. We combined RNA-Seq-based transcriptomics with bioinformatical, biophysical and behavioural tools to explore the landscape of age-variable genes of Johnston’s Organ (JO) - the flies’ inner ear’. Our data suggests that the thereby identified transcriptional regulators are not restricted to *Drosophila* - or the sense of hearing - but represent key players of homeostasis across taxa and probably across sensory modalities.

## Results

### *Drosophila* is prone to age-related hearing loss (ARHL)

Functionally, the *Drosophila* antennal ear (Fig. 1a) is composed of two components: (i) the external *sound receiver* (jointly formed by the third antennal segment, A3, and its lateral appendage, the arista) and (ii) the actual *‘inner ear’*, which is formed by Johnston’s Organ (JO), a chordotonal organ ^22^ located in the second antennal segment, A2. JO harbours ~500 mechanosensory neurons ^23^.

**Figure 1.**
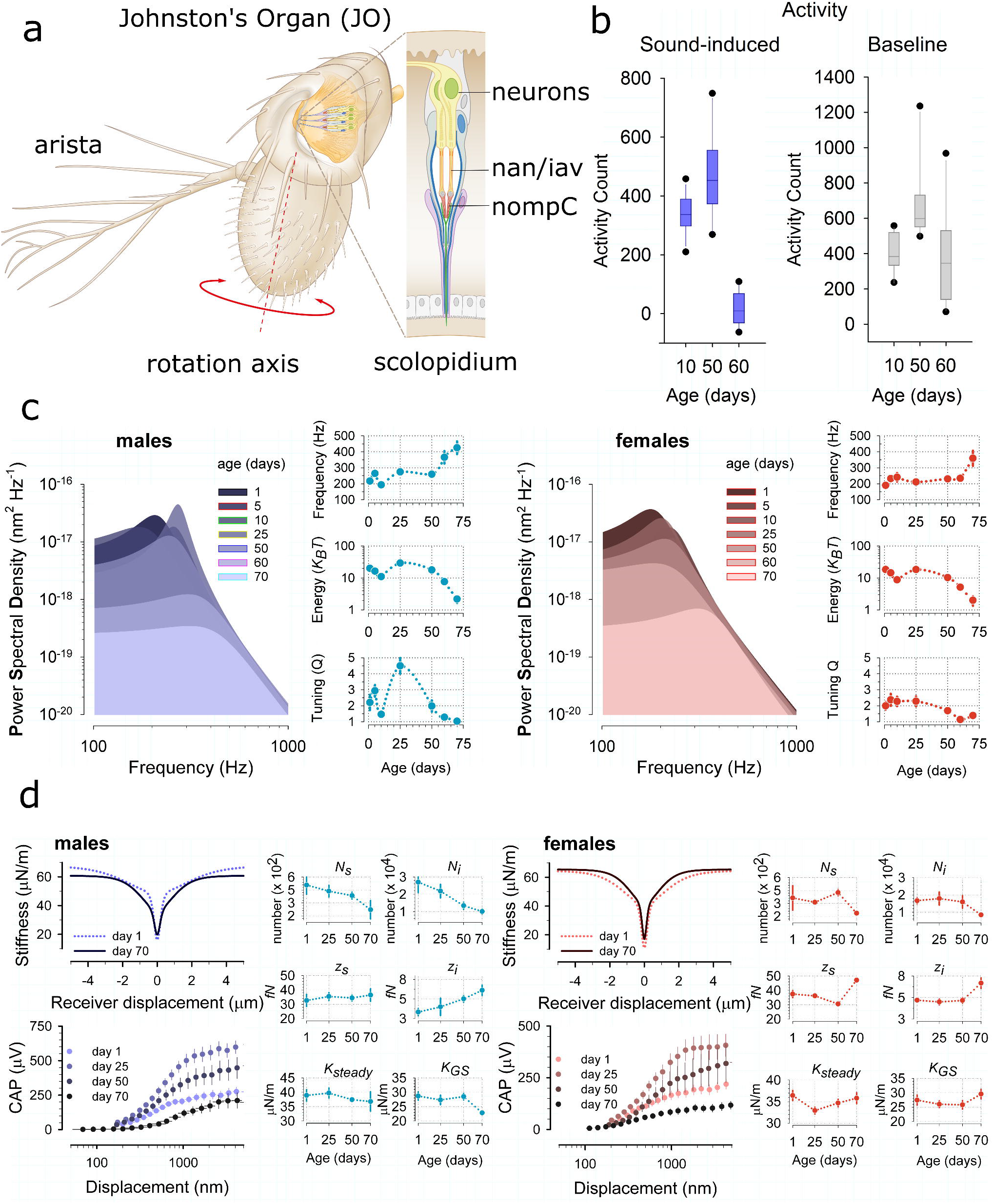
*Drosophila* Hearing across the life course. (a) Schematic representation of Johnston’s Organ (JO), a chordotonal organ located in the 2^nd^ antennal segment. JO harbours the mechanosensory units (scolopidia) that mediate the sensation of sound in *Drosophila*. Sound waves act on the feathery arista, forcing the 3^rd^ antennal segment to rotate about its longitudinal axis, thereby stretch-activating specialised mechanosensory ion channels (Nan, Iav, NompC) in the scolopidial neurons. (b) Sound-evoked activity (left, male locomotor responses to courtship song) is abolished in 60 day old flies (p=1.004E-012, t-test). Baseline activity levels (right, male locomotor activity when not stimulated) are not significantly different between 10 and 60 day old flies (p=0.487, t-test). (c) Power Spectral Densities of unstimulated antennal sound receivers betray age-related decline of hearing in both males (left, shades of blue) and females (right, shades of red). Preceded by homeostatic oscillations around their baseline values, all principal parameters of hearing (shown in right-hand panels for both sexes) indicate a loss of hearing from day ~50 onwards: the receiver’s best frequency starts rising towards the level of the passive system, the auditory energy gain drops to near zero and tuning sharpness falls to values around ~1. (d) Mechanical and electrophysiological responses to force steps allowed for probing JO mechanotransducer function across the auditory life course in male (left, blue) and female (right, red) flies. Mechanical integrity of auditory transducers was quantified by fitting gating spring models to the antennal receiver’s dynamic stiffness (slope stiffness) as a function of its peak displacement (see ref. ^13^ for details). Electrophysiological function was assessed by recording compound action potential (CAP) responses from the antennal nerve. CAP responses showed an identical pattern across the life course in both males and females: CAP response magnitudes substantially increased from day 1 to day 25, then monotonously declined from day 25 to day 70. The largest drop in CAP magnitudes occurred between day 50 and day 70, with responses of 70-day-old flies even falling below those of 1-day-old flies. Transducer mechanics, in contrast, remained more intact throughout. However, at day 70 the five principal parameters of transducer function, i.e. the number of sensitive transducer channels *N*_*s*_), the number of insensitive transducer channels (*N*_*i*_), the sensitive single channel gating force (*z*_*s*_), the insensitive single channel gating force (*z*_*i*_) and the total gating spring stiffness (*K*_*GS*_) were all significantly different from their values at day 1, in both males and females. Interestingly, no such change was observed for the stiffness of the antennal joint (*K*_*steady*_), which is a transducer-independent measure of antennal mechanics. Next to these properties shared between males and females, our analyses also revealed some sexually dimorphic phenomena: *K*_*GS*_ goes up in 70-day-old females, but it goes down in 70-day-old males. Whereas in females *N*_*s*_, *N*_*i*_, *z*_*s*_ and *z*_*i*_, remain at constant values until the age of 50 days, the respective values of male flies change monotonously throughout the life course, with continually falling numbers of transducer channels being compensated by increasing single channel gating forces (thereby homeostatically balancing the male antenna’s nonlinear stiffness).

To assess hearing across the *Drosophila* life course we first measured the locomotor activities of flies in response to a playback of courtship song components at different ages. *Drosophila melanogaster* males increase locomotor activity in response to courtship song ^24^. While 10- and 50-day-old flies increased their locomotor activity in response to a 15 min long train of courtship song pulses (inter-pulse-interval, IPI: 40ms), sound-induced responses were absent in 60-day-old flies (Fig. 1b, left). Baseline locomotor activities of 60-day-old flies, however, remained the same as in 10-day-old flies (Fig. 1b, right), suggesting an auditory - rather than a more generalised neurological - deficit as the underlying cause for the non-responsiveness to sound.

A simple, but quantitatively powerful, test of auditory performance was then conducted by recording the vibrations of unstimulated sound receivers (*free fluctuations*) ^15^. A receiver’s free fluctuations reveal three principal parameters of auditory function: (1) the ear’s best frequency, *f*_*0*_ (measured in Hz), (2) its frequency selectivity or quality factor, *Q* (dimensionless), and (iii) its energy - or power - gain (measured in *K*_*B*_*T*). Much like hair cells in the vertebrate inner ear, the antennal ears of *Drosophila* are active sensors, which inject energy into sound-induced receiver motion ^25^.

Our data show that the ears of flies, much like those of humans, show age-related hearing loss (ARHL) (Fig. 1c). At 25 °C, the antennal receivers of 70-day-old flies show (i) best frequency shifts towards the passive system, where no energy injection is observed, (ii) a greatly reduced tuning sharpness and (iii) a ~90% loss of their energy gains (Fig. 1c and Supplementary Table 1), indicating a near-complete breakdown of the active process - which supports hearing - at day 70. The time course of this auditory decline was broadly similar between males and females (Supplementary Table 1).

To probe auditory function in more detail, we also quantified the mechanical and electrophysiological signatures of auditory mechanotransduction in response to force-step actuation of the fly’s antennal ear at different ages (Fig. 1d). Direct mechanotransducer gating introduces characteristic nonlinearities - namely drops in stiffness - into the mechanics of the sound receiver. These so-called ‘gating compliances’ can be modelled with a simple gating spring model ^13,26^ thereby allowing for calculating the number - and molecular properties - of different populations of mechanosensory ion channels present in the fly’s JO ^27^. Two distinct mechanotransducer populations have previously been described: a *sensitive* population, linked to hearing, and an *insensitive* population, linked to the sensation of wind and gravity ^16^. At day 70, the numbers of predicted sensitive (*N*_*s*_) and insensitive (*N*_*i*_) channels have decreased by ~50% as compared to their values at day 1; the single channel gating forces of the sensitive (*z*_*s*_) and insensitive channels (*z*_*i*_), in turn, have increased (Fig. 1d). The receiver’s steady-state stiffness (*K*_*steady*_), however, which is an indicator of the integrity of the antennal joint, is not significantly different between 1- and 70-day-old flies, suggesting that the changes in auditory mechanics reflect an ageing of the mechanotransducer machinery rather than structural changes of the organ itself. Parallel recordings of compound action potentials (CAP) from the antennal nerve showed that nerve response magnitudes initially increased from day 1 to day 25 and then decreased steadily, with response curves of 70-day-old flies falling below those of 1-day-old flies, both in response magnitude and displacement sensitivity (Fig. 1d). The above-described pattern of transducer ageing was seen in both males and females. Some subtle differences, however, could be observed between the sexes. While females displayed a ~stable baseline of most transduction parameters up to day 50, males showed signs of a more steady decline from day 1 on. Also, gating spring stiffnesses (*K*_*GS*_) decreased in 70-day-old males but increased in 70-day-old females (Fig. 1d).

Summing up the behavioural, mechanical and electrophysiological evidence, the auditory life course of *Drosophila melanogaster* can roughly be broken down into two phases: (i) a dynamic phase of *homeostatic metastability*, which is characterised by fluctuations of key parameters of hearing around a ~stable baseline, which last from day 1 to ~day 50 (also including possible signs of initial functional maturation) and (ii) a phase of *terminal decline*, which starts at ~day 50 and leads to a near complete loss of auditory function at ~day 70.

We hypothesized that a breakdown of the homeostatic machinery that shapes auditory performance during the life course - and maintains healthy hearing up until day 50 – is the ultimate reason for the observed terminal decline. In order to identify the molecular networks involved, we therefore profiled the auditory transcriptome at days 1, 5, 10, 25 and 50 through RNA sequencing (RNA-Seq) of the 2nd antennal segment (Supplementary Table 2).

### The age-variable auditory transcriptome in *Drosophila*

16,243 genes are expressed in the 2^nd^ antennal segment in both males and females (Supplementary Table 2); 13,324 of those are protein-coding. We compared the expression levels of all genes in a pair-wise manner, between (i) day 1 and 5, (ii) day 5 and 25 and (iii) day 25 to 50. In total, 5,855 (4,936 protein-coding) genes were changing their expression significantly in at least in one of the three pair-wise comparisons (criteria: >1.5-fold change; <10% False Discovery Rate (FDR); p<0.05; Supplementary Table 3 and Supplemental Methods).

The gene-ontological nature of age-variable genes in A2 was probed with the **G**ene **O**ntology en**RI**chment ana**L**ysis and visualization (GOrilla) tool ^28,29^. The age-variable transcriptome revealed both down- and upregulation of genes. Genes involved in ATP metabolism, protein processing and structural molecules were found to be downregulated, whereas immune response genes, photo transduction genes and translation machinery genes were upregulated (Fig. 2a and Supplementary Table 4). Next to many novel JO genes, about one third (109) of all previously reported JO genes (314) ^19,21^ changed their expression in our dataset (Table 1); this included rhodopsins, the TRPV channel gene *nan*, innexins, as well as ATPase β subunits (nervanas) previously linked to JO function ^30,31^.

**Figure 2.**
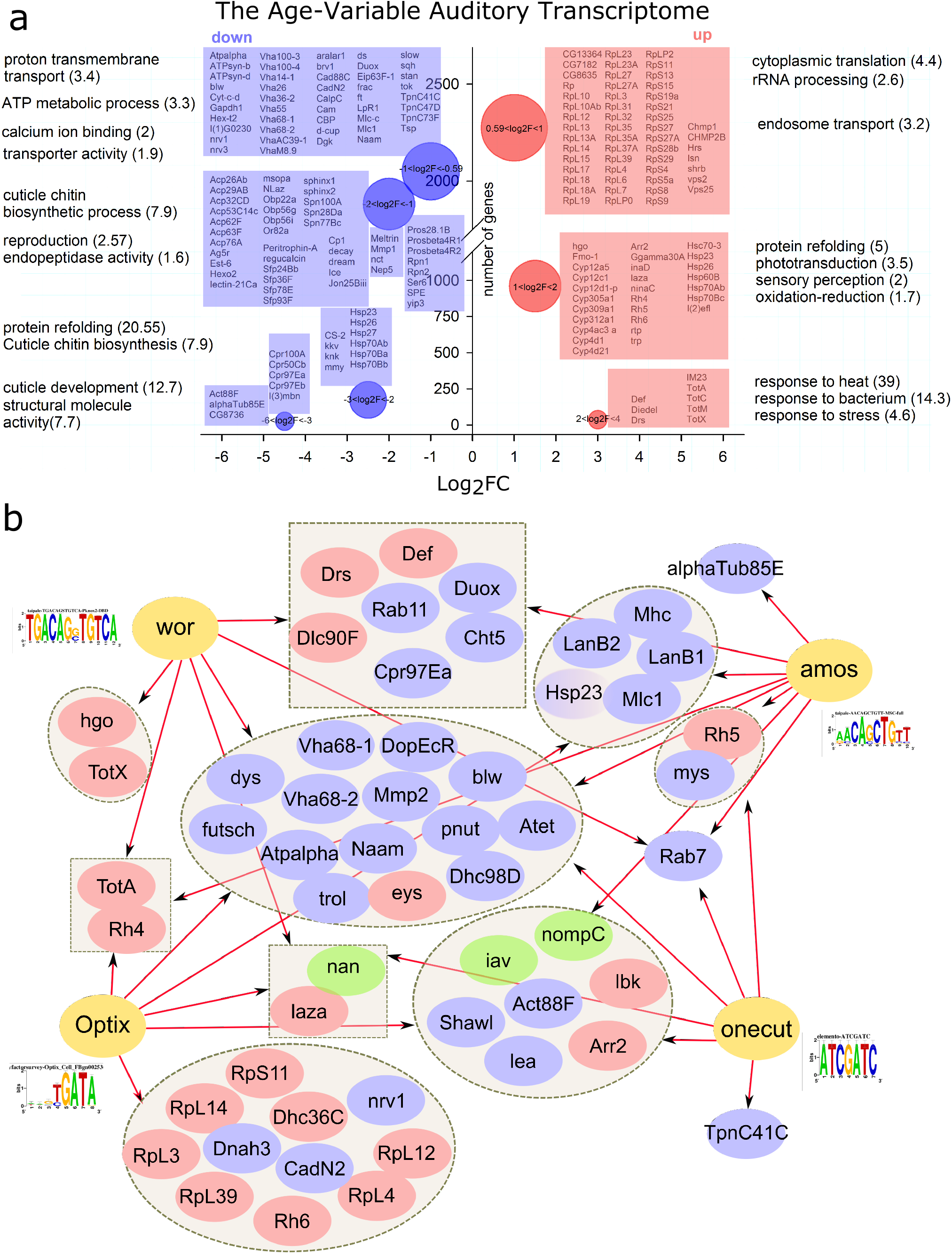
Gene-Ontology and Bioinformatics of the *Drosophila* age-variable JO transcriptome. (a) Gene Ontology (GO) based summary of age-variable genes in JO as derived from RNAseq data taken across different age points (days 1, 5, 10, 25 and 50). Down-regulated (blue) and up-regulated (red) genes from multiple pairwise comparisons between all age points are shown on the left and right side of the graph, respectively. The bubble diameter is proportional to the gene number (the larger the diameter the more genes were eon-or up-regulated.The y-axis shows numbers of the genes and the x-axis the Log_2_Fold-Change (Log_2_FC) of gene expression. GO terms correspond to the up-regulated (right) or down-regulated (left) genes, a selection of which is shown in the respective neighbouring boxes; enrichment scores are shown in brackets. Selection of the most age variable genes are shown in individual boxes corresponding to each GO term next to it. (b) Prediction of upstream transcriptional master regulators to age-variable JO target genes (based on motif-binding analysis in the *i*Regulon software package). Identified master regulators *wor, amos, Optix* and *onecut* are shown in yellow with arrows leading to their predicted targets. Targets are grouped and up and down-regulated genes are shown in blue and red, respectively. Mechanosensory ion channels, previously linked to fly hearing, are shown in green: *iav* and *nompC* are predicted to be downstream of *onecut*, *amos* and *Optix*, whereas *nan* is predicted to be downstream of *wor*, *Optix* and *onecut*.

**Table 1.**
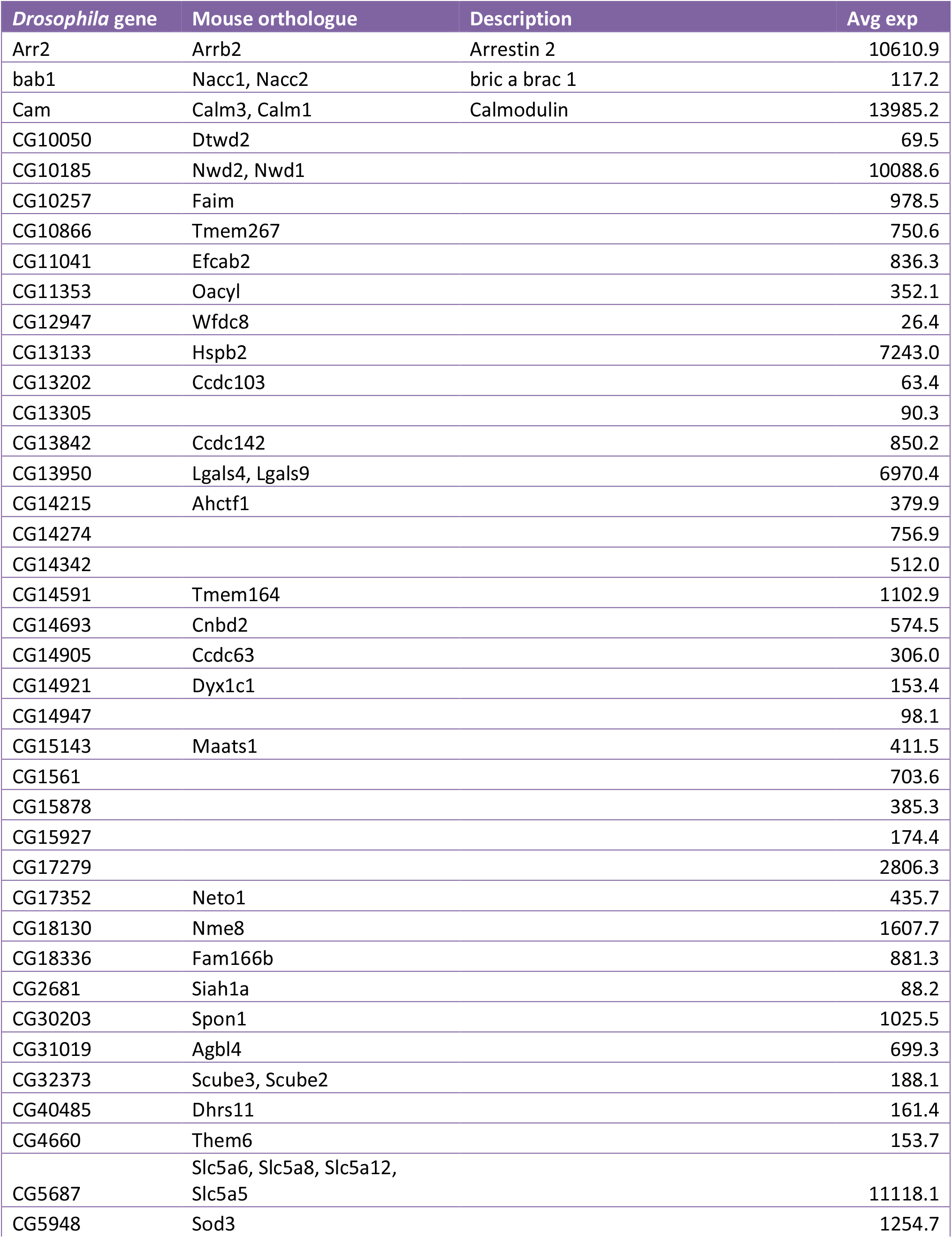
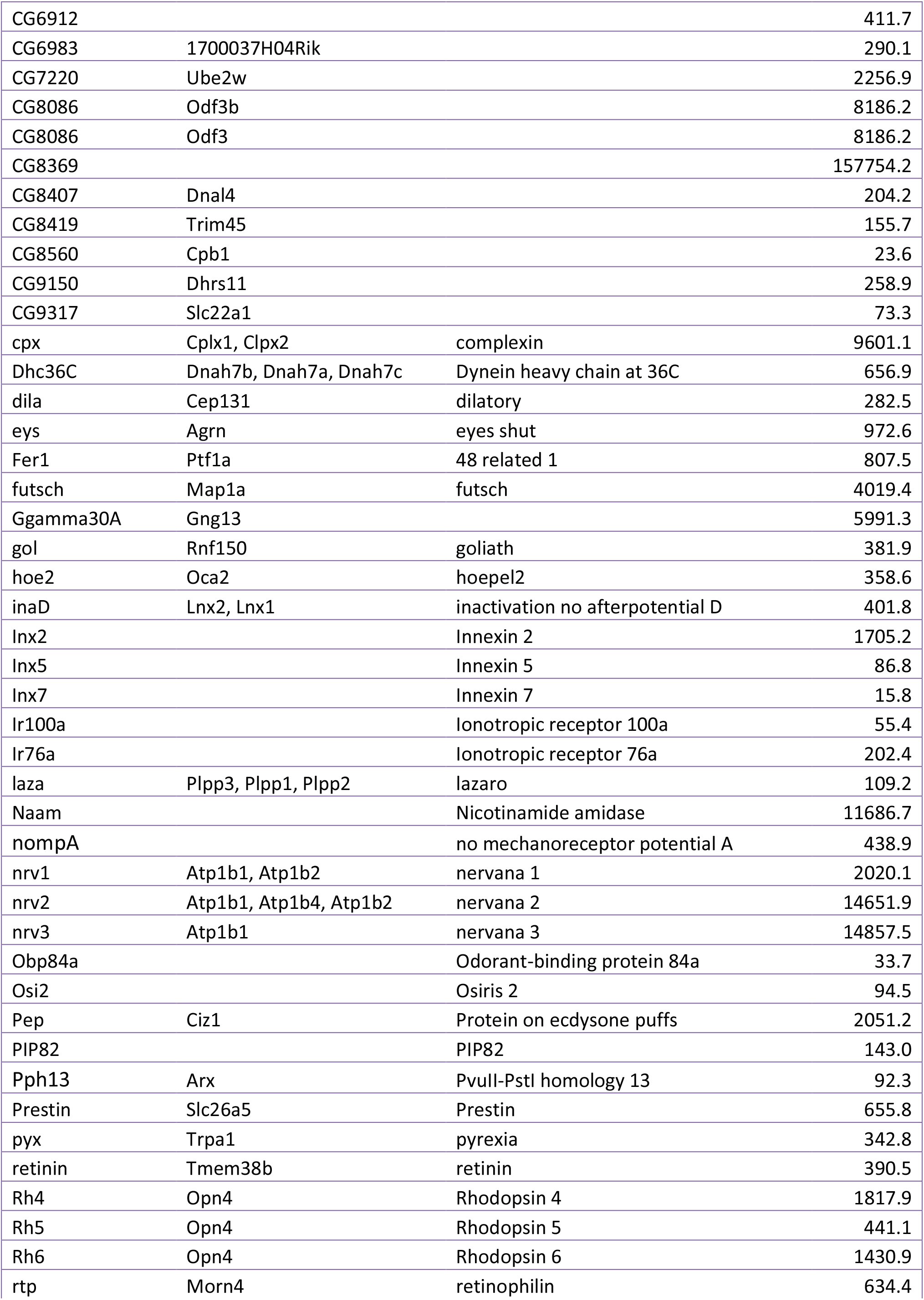
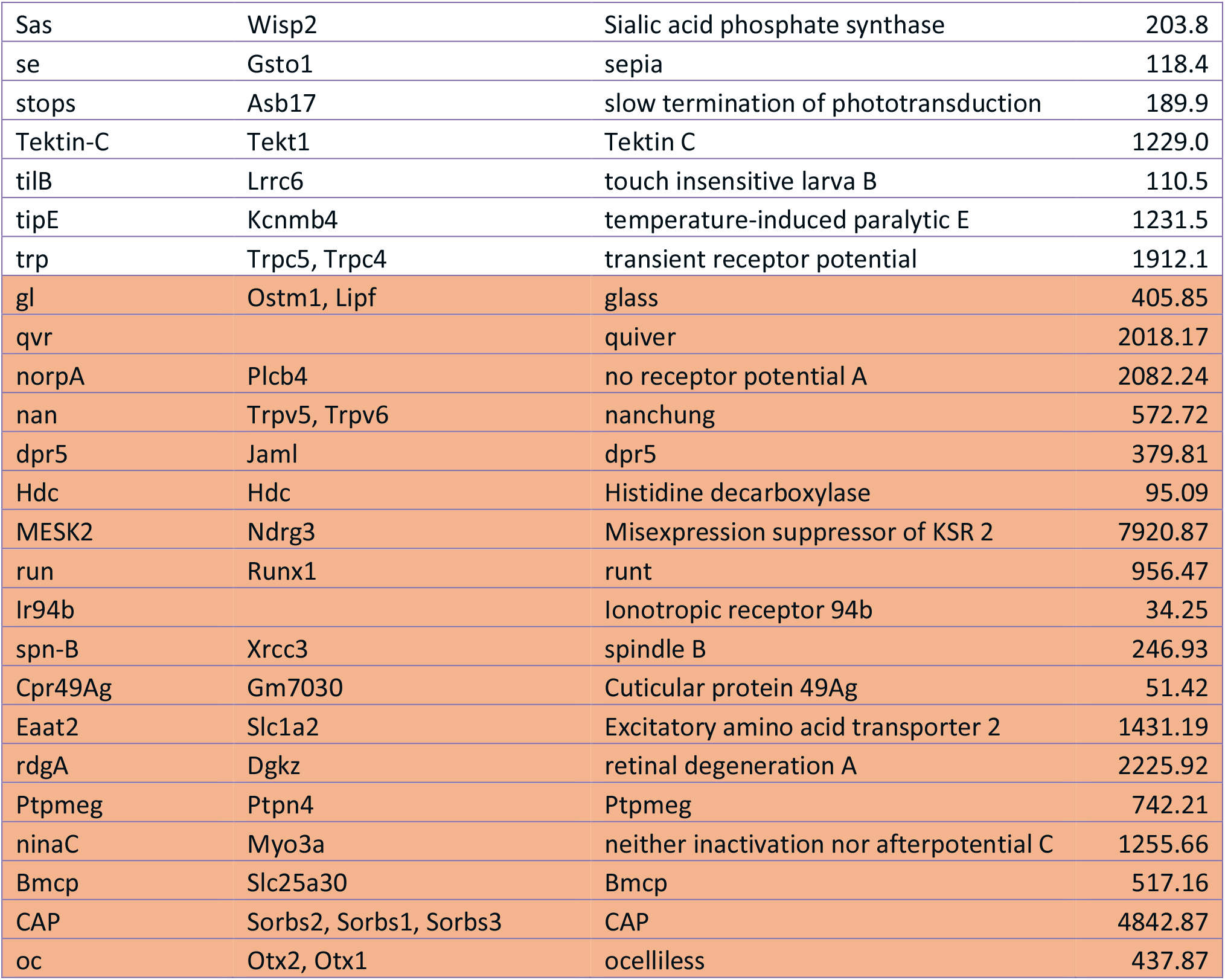
Previously identified JO genes showing age-variable expression. Highlighted genes in orange change their expression only in males. Avg exp stands for Average expression

We also found that 67% (74 out of 111) of recently identified mammalian hearing loss genes ^10–12^ are conserved in flies - and expressed in A2 – with 32% of them also showing age-variable expression in JO (Table 2).

**Table 2.**
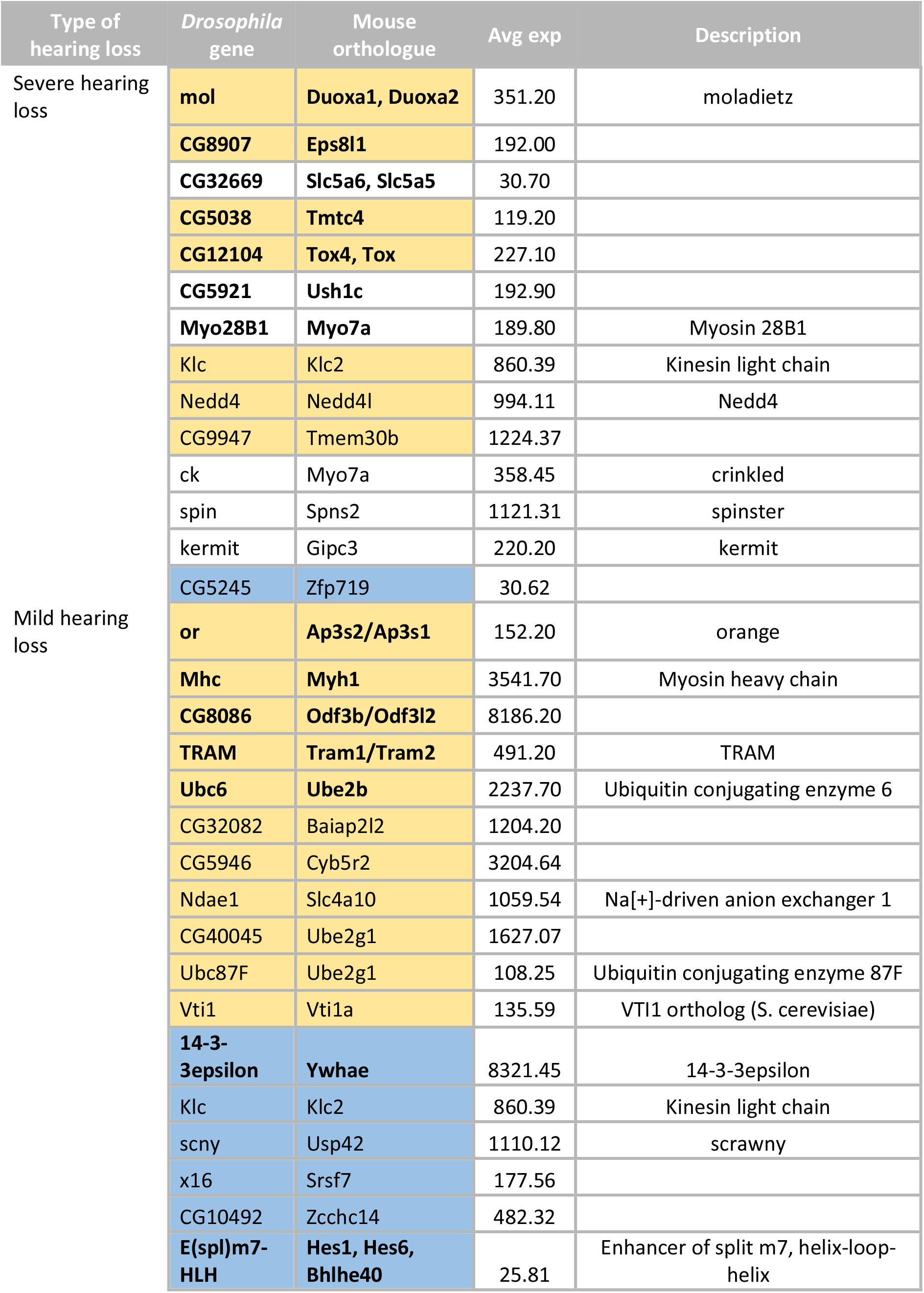
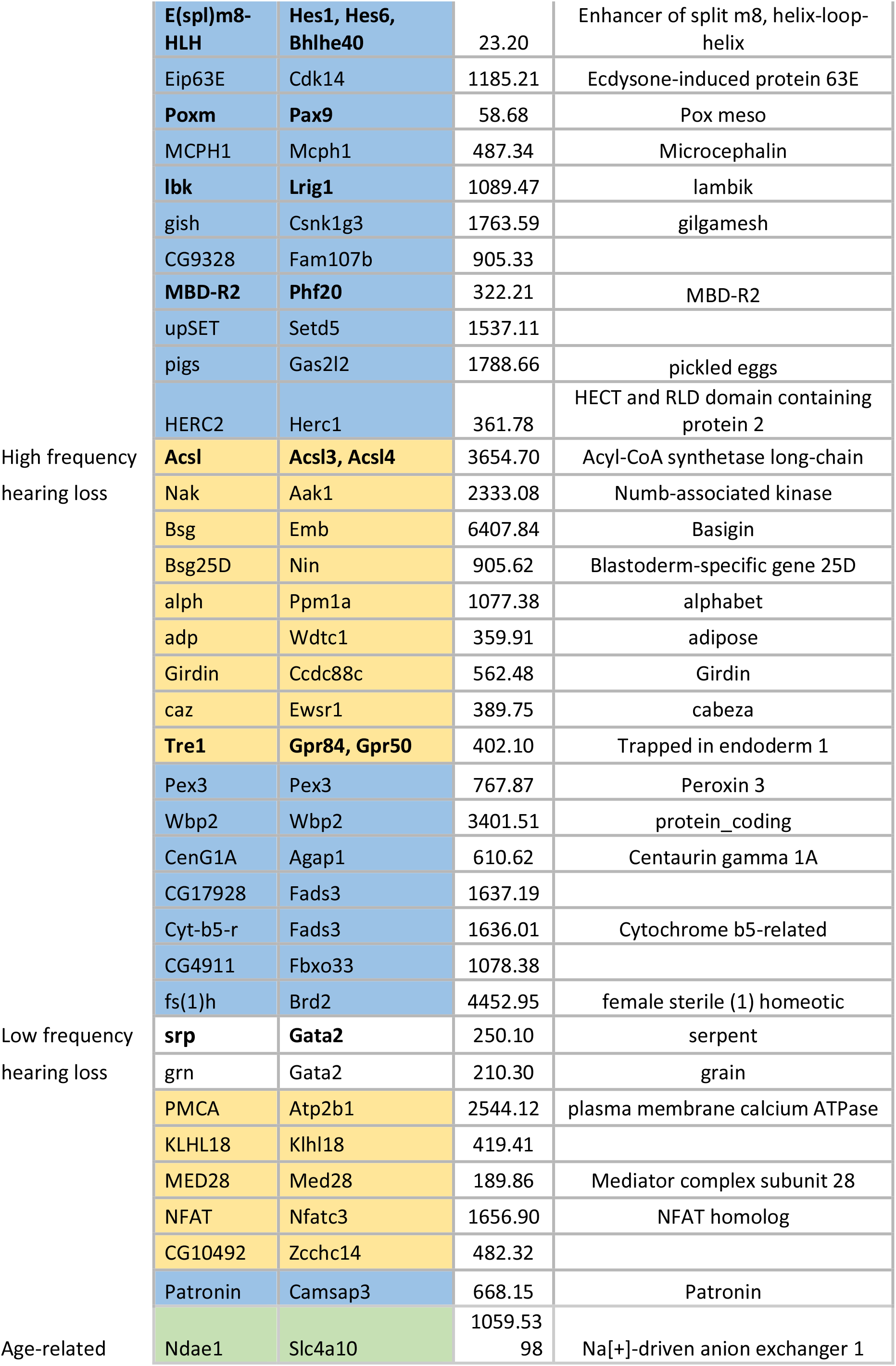
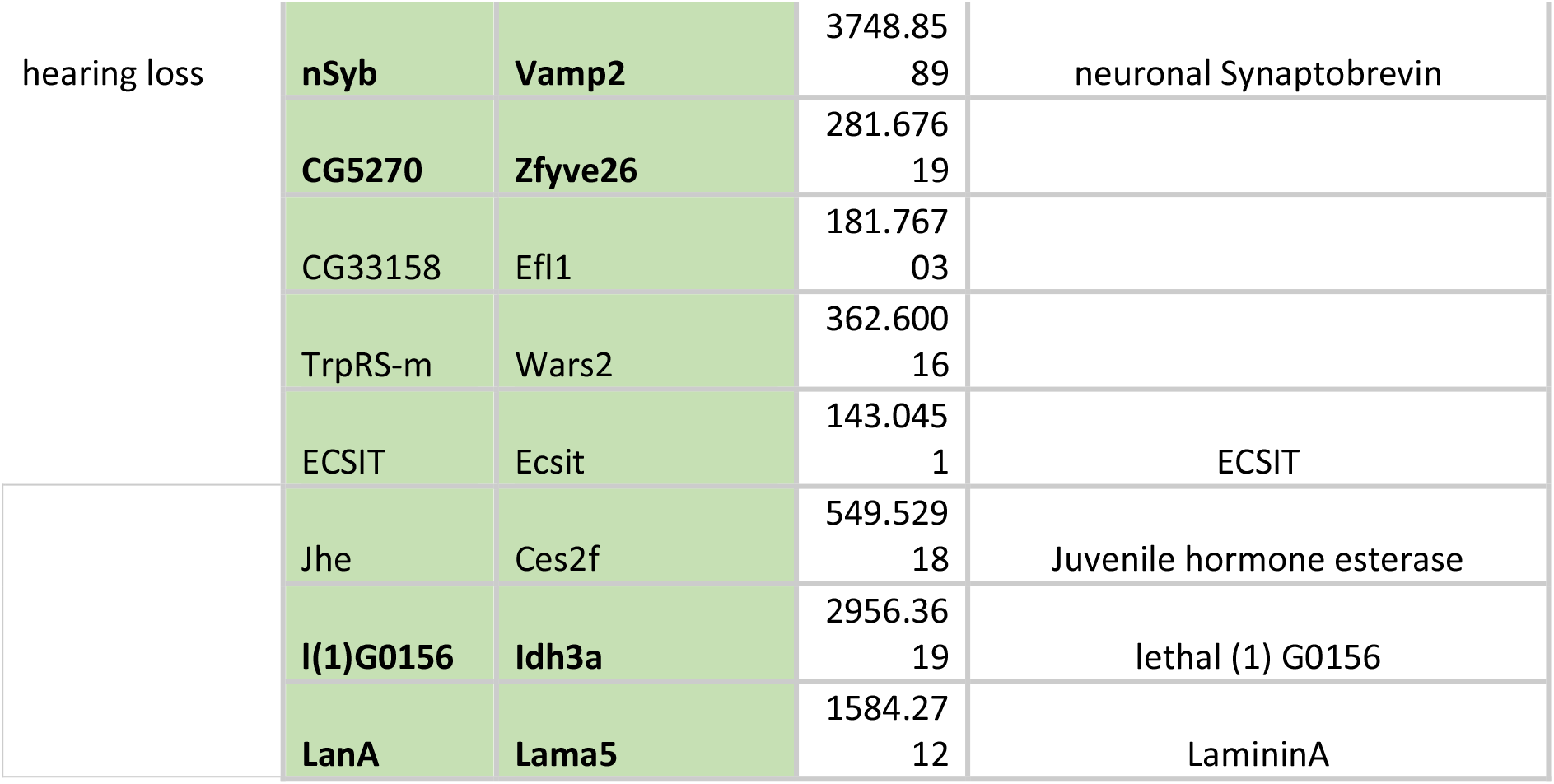
Mouse genes linked to deafness are conserved and expressed in fly JO. Bowl R et al, 2017 genes are shown in light orange, Ingham N et al 2019 genes are shown in blue and Potter P et al 2016 age-related genes are shown in green. JO age-variable genes are shown in bold.

### Number of JO neurons remains constant up until the age of 50 days

To test whether the age-variable transcriptome between day 1 and day 50 reflected changes on the cellular level we counted the number of neurons in the second antennal segment at different ages (Supplementary Fig. 1). From day 1 to day 50 no difference in neuronal numbers was seen, suggesting that the observed transcriptomic changes betray an age-variable transcriptional - i.e. gene-regulatory - activity.

### Predicting the gene-regulatory landscape of auditory homeostasis in *Drosophila*

In order to shed light on the gene-regulatory networks of auditory homeostasis and identify key transcription factors (TFs) acting upstream of the age-variable genes, we applied the bioinformatics software package iRegulon, which predicts TFs based on motif binding probabilities ^32^.

Our heuristic rationale was based on three assumptions: (i) auditory homeostasis involves, at least in parts, specific TFs; (ii) TFs can be low abundance genes, their action can be mediated through small changes in expression levels; (iii) every TF has, on average, more than one target and those targets might be functionally, or gene-ontologically, linked.

We drew three consequences from the above premises: First, we concentrated our study on TFs. Second, we used the entire age-variable auditory regulon to *predict* upstream TFs, thereby increasing the overall sensitivity of our analysis. Even TFs, which might have escaped our attention from the RNA-Seq data itself could thus be recovered in subsequent bioinformatical analyses. Third, we grouped putative regulons (i.e. subsets of expressed genes) not only by their variability with age but also by their gene-ontological classification.

37 TFs were predicted from different rounds of gene submission (Supplementary Table 5), based on varying gene ontological categories, such as (i) transporters and receptors, (ii) trafficking genes, (ii) structural genes, (iv) most abundantly expressed genes or (v) genes most variable between ages (Fig. 2b and Supplementary Fig. 2).

### Testing predicted homeostatic regulators of *Drosophila* hearing

To test the validity, and functional relevance, of the bioinformatical analyses, we used RNAi-mediated, adult-specific knockdowns (KDs) of 19 (out of 37) predicted transcription factors: *Adult enhancer factor 1* (*Aef1*), *absent MD neurons and olfactory sensilla* (*amos*), *anterior open* (*aop*), *araucan (ara)*, *atonal* (*ato*), cut (*ct*)*, glass* (*gl*)*, longitudinals lacking* (*lola*)*, onecut, Optix, pannier* (*pnr*)*, PvuII-PstI homology 13* (*Pph13*)*, regulatory factor X (Rfx)*, *runt (run), Sox box protein 14* (*sox14*)*, serpent* (*srp*)*, Signal-transducer and activator of transcription protein at 92E* (*Stat92E*)*, TATA-binding protein* (*Tbp*) and *worniu (wor)* (Supplementary Table 6). Analysing the free fluctuations of the antennal sound receiver, we found 5 cases (*onecut*, *amos*, *gl*, *lola* and *Sox14*), where the knockdown accelerated the ARHL phenotype; 4 other cases (*wor*, Optix, *Pph13* and *ara*), however, showed protective phenotypes in various principal parameters of auditory function (Supplementary Table 6). To our surprise, adult-specific knockdowns of the crucial developmental genes *ato* ^18^, *Rfx* ^33^ and *ct ^34^* did not show any significant phenotypic changes (Supplementary Table 6), suggesting that they are not involved in homeostatic maintenance of hearing in adults.

To get a better understanding of the specific TF-mediated homeostatic programme that maintains hearing, we concentrated on the top four regulators, which occurred consistently throughout various rounds of bioinformatical analyses. These were *onecut*, *Optix*, *wor* and *amos*, all of which showed clear expression in the neurons of JO (Fig. 3). These four TFs also showed the strongest KD phenotypes in the free fluctuation experiments (Fig. 2b, Supplementary Table 6 and Fig. 4a, b), with each TF affecting distinct aspects of auditory function. Analysing the mechanical and electrophysiological signatures of mechanotransducer gating across the four KDs (Fig. 4c, d) identified *onecut* as a crucial homeostatic regulator of auditory transducer function. The number of predicted sensitive (auditory) transducer channels (*N*_*s*_) is greatly reduced in *onecut* KD flies, while their single channel gating forces (*z*_*s*_) are increased. The numbers of predicted insensitive (non-auditory) channels (*N*_*i*_) are slightly increased and their single channel gating forces (*z*_*i*_) decreased in *onecut* KDs. The observed inverse relationship between ion channel numbers and gating forces might represent an intrinsic homeostatic link between the two parameters (see also discussion and Supplementary Fig. 3). CAP responses to force-step actuation, finally, are dramatically reduced in the KD condition. The overall effect of the adult-specific KD of *onecut* is a near-complete abolition of the mechanical and electrical signatures of sensitive auditory transducer gating. Consistent with these results, *onecut* KD flies specifically lose their responsiveness to sound, while their baseline locomotor activities remain unchanged (Fig. 4e). KDs of *Optix*, *amos* and *wor* showed less pronounced effects on electrophysiological or mechanical signatures of transducer gating, but at least one transducer parameter was affected in each genotype (Fig. 4d). For three of the four master regulators (Optix, wor, amos), overexpression constructs were available, we thus also explored whether overexpression could invert the knockdown phenotypes seen in the free fluctuation analyses (compare to Fig. 4a); this was indeed the case for *Optix* and *amos*; wor overexpression was indistinguishable from the controls (Supplementary Fig. 4, Supplementary Table 6). Over-expression of *amos* and downregulation of *wor* for 30 days at 30 °C led to a partial prevention of the age-related auditory decay (Fig. 5).

**Figure 3.**
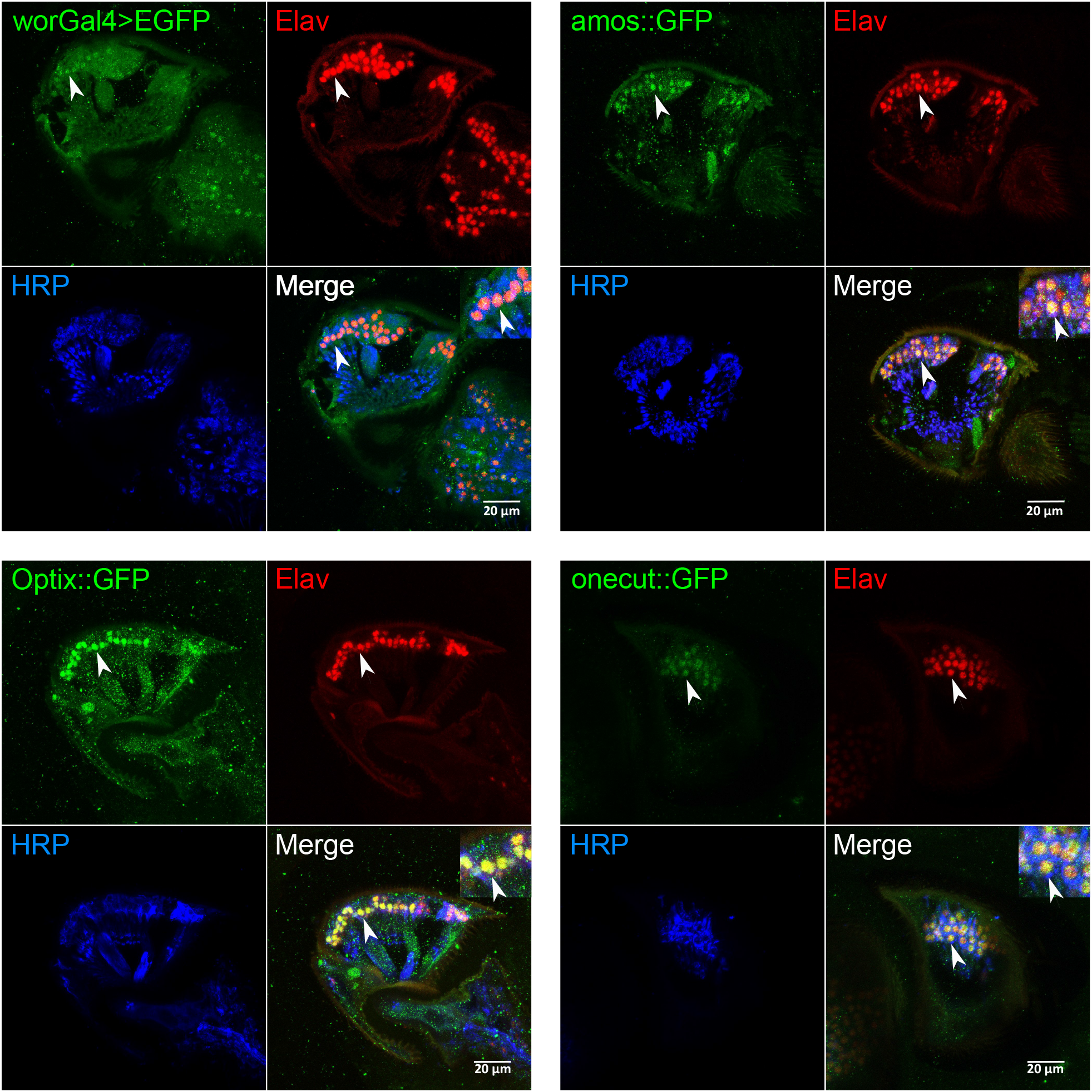
Expression validation of homeostatic master regulators in JO. All four predicted regulators (Wor, Amos, Optix and Onecut) are expressed in JO (expression analysis was done at the age of day 10 for all genotypes). Expression of Wor was detected by expressing EGFP under the control of a *wor*-Gal4 driver; expression of Amos, Optix and Onecut was detected by using GFP-tagged flyFos gene expression constructs ^58^. Co-labelling with antibodies against two pan-neuronal markers (the nuclear marker Elav, red; and the membrane marker HRP, blue) confirmed neuronal expression for all four regulators. Arrowheads indicate examples of clear co-localization between the three signals.

**Figure 4.**
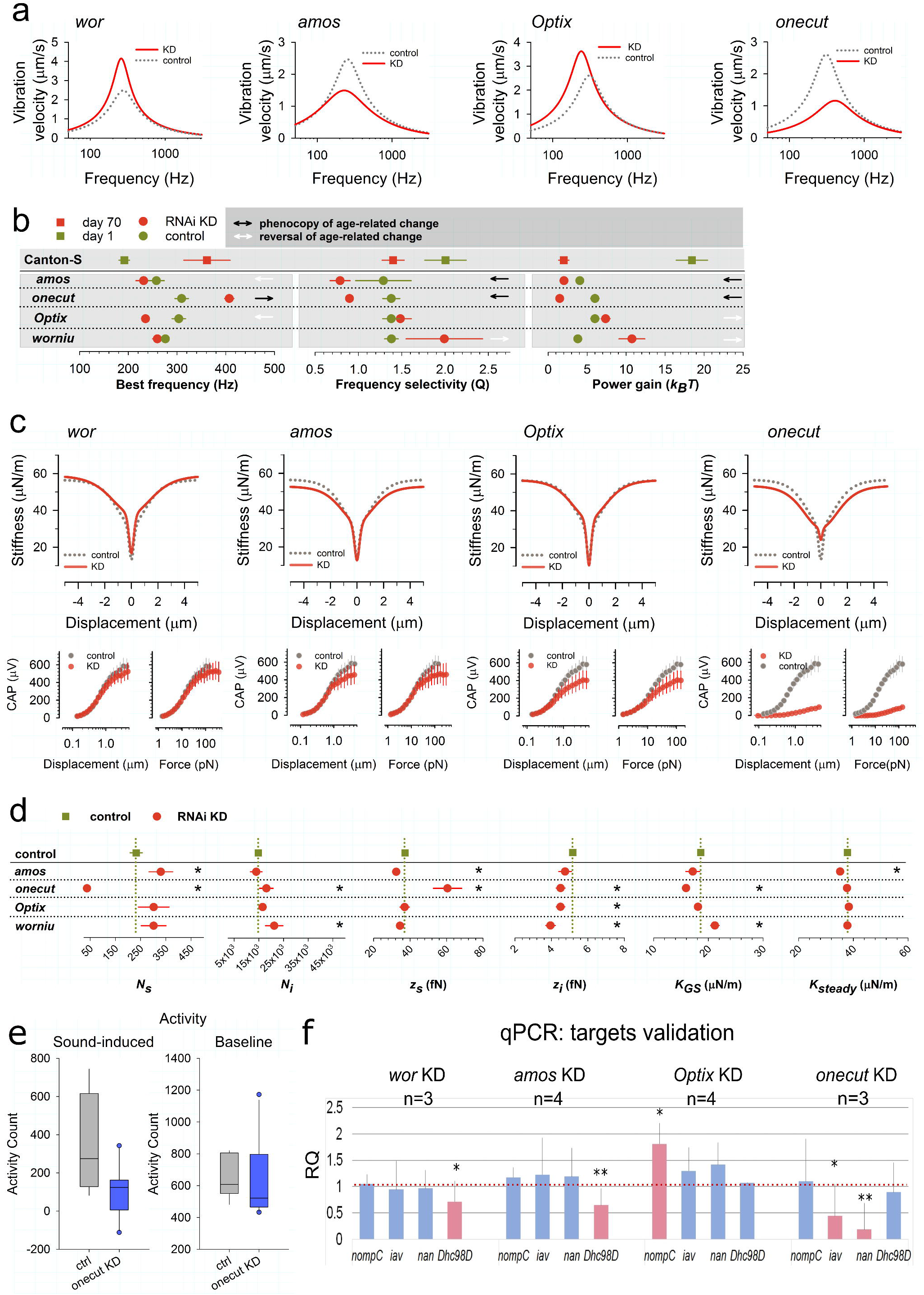
Functional validation of homeostatic master regulators. (a) Average vibration velocities of female unstimulated sound receivers (‘free fluctuations’) after adult-specific, RNAi-mediated knockdown (KD; red solid lines) for all four master regulators alongside their respective controls (grey dashed lines). KDs of *amos* and *onecut* show a loss of sound receiver function, as evident from (i) reduced energy content (‘power gain’), (ii) reduced frequency selectivities, and - in the case of *onecut* - also (iv) best frequency shifts towards higher values. KDs of *wor* and *Optix*, in contrast, show enhanced sound receiver function, as evident from (i) increased energy content and (ii) increased frequency selectivity (*wor*) or best frequency shifts to lower values (*Optix*). [Supplementary Table 6 for numerical details and statistics] (b) Line plot summaries comparing the KD sound receiver phenotypes [as from (a)] to the sound receiver phenotypes occurring naturally during ageing (reference for comparison: Canton-S day 1 to day 70). Arrows indicate significant changes in parameters. Black arrows indicate that KD phenotypes (relative to their corresponding controls) phenocopy the age-related hearing loss (ARHL) phenotypes seen in wildtype flies. White arrows indicate a reversal of the specific ARHL phenotype. (c) Gating compliances (average fits, top) and CAP responses (medians plus standard errors, bottom) to force step actuation across adult-specific KDs of four master regulators (red) and their corresponding controls (grey). CAP responses are plotted against both stimulus force and antennal displacements. KD of *onecut* leads to a dramatic loss of auditory transducer function, as evident from the near complete loss of the gating compliance for the most sensitive transducers and the loss of nerve responses to small stimulus forces/displacements. KDs of *wor*, *amos* and *Optix* have subtler effects on transducer mechanics but all reduce nerve responses to larger stimulus forces/displacements. (d) Line plot summaries of transducer mechanics [from (C)] in four regulator KDs (red) relative to controls (green). Dashed lines indicate respective control values. Significant changes are asterisked (*). (e) Sound-induced behavioural responses in males after *onecut* KD (blue) compared to control flies (grey). *onecut* KD mutants do not respond to sound (left, p=0.0127, t-test), while their baseline activities are unaltered (p=0.351, Mann-Whitney Rank Sum Test). (f) Gene expression changes after regulator KDs as quantified by RT-qPCR. *wor* and *amos* KDs show significant reduction of the dynein motor Dhc98D, while KD of *Optix* leads to overexpression of NompC; *onecut* KD reduces expression of both *nan* and *iav*. (*n* indicates the biological replicates, error bars show standard deviations, *p> 0.05, **p>0.01).

**Figure 5.**
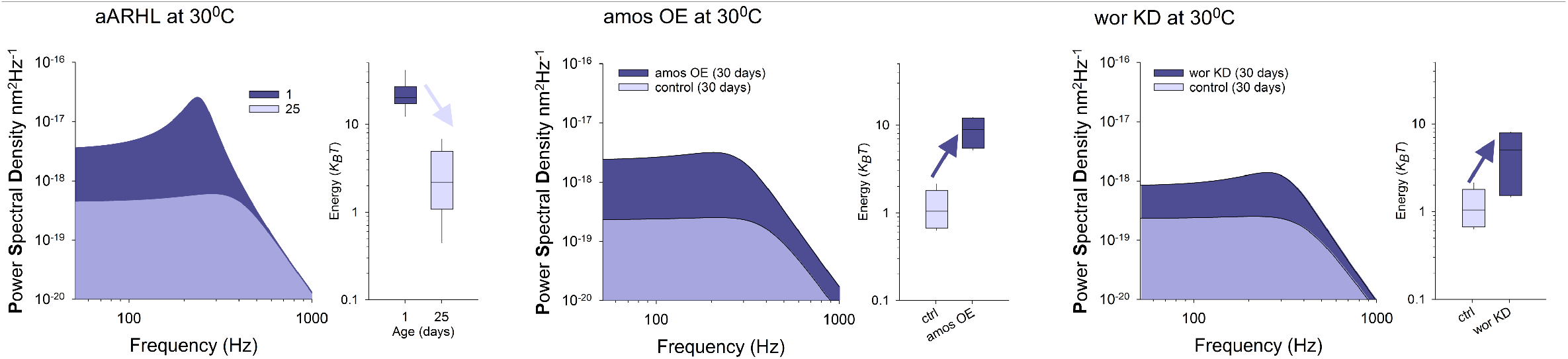
*amos* overexpression or *wor* knockdown reverse the course of hearing loss and enhance auditory function. (left) Power Spectral Densities of unstimulated antennal sound receivers betray accelerated age-related hearing loss (aARHL) in flies kept at 30 °C (left), with a near complete loss of receiver activity already at ~day 25 (light blue area: 1 day old flies; dark blue area: 25 day old flies). A 30-day-long *amos* (middle) overexpression (OE) or *wor* (right) knockdown (at 30 °C) protects receivers from the age-related loss of activity (dark blue: KD or OE, respectively; light blue: controls). Box plots show energy contents (power gains) for each transgenic intervention (dark blue) relative to controls (light blue).

### qPCR validation reveals key auditory targets of master regulators

Knockdown and overexpression of identified homeostatic TFs altered important parameters of the fly’s ear, such as its frequency tuning, mechanotransduction, amplification and nerve responses (Fig. 4a-d). All of these system properties are thought to arise from an interaction of three key transient receptor potential (TRP) channels, namely Nanchung (Nan) ^35^, Inactive (Iav) ^36^ and NompC ^37^ with motor proteins from the dynein family ^20^. One such dynein was also identified within our age-variable gene set, this is the Dynein heavy chain at 98D (Dhc98D). The three TRP channels from above, as well as the auditory dyneins were predicted downstream of the four master regulators (Fig. 2b, Supplementary Fig. 2). Using real-time quantitative polymerase chain reactions (qPCRs) we therefore tested if *nan*, *iav*, *nompC* and *Dhc98D* levels were under the control of the identified homeostatic TFs (Fig. 4 f). RNAi-mediated adult-specific knockdown of *onecut* resulted in a dramatic downregulation of both *nan* and *iav*, knockdown of *Optix* lead to an upregulation of *nompC* levels, whereas knockdown of *amos* and *wor* showed downregulation of *Dhc98D*.

Adult-specific knockdown of *Dhc98D* caused a strong hearing loss phenotype similar to the one seen after *amos* KD (Fig. 6 and Fig. 4a).

**Figure 6.**
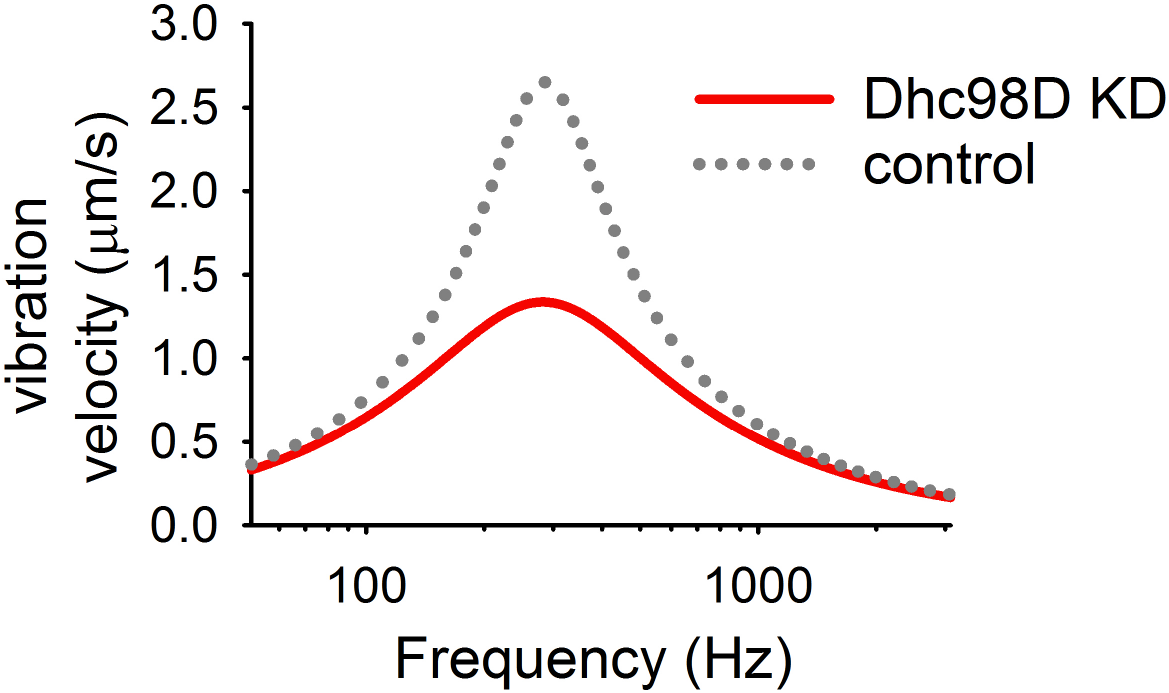
*Dhc98D* gene adult-specific knockdown (KD) Vibration velocity of the sound-receiver and the sharpness of the tuning Q are significantly reduced in knockdown flies (shown in red) compared to the controls (shown in dotted grey). See also Supplementary Table 6.

## Discussion

We show that the ears of flies, just like those of humans, are prone to age-related hearing loss (ARHL). ARHL manifests in all aspects of hearing function. More surprising than its eventual decay, however, is the long period during which sensitive hearing is preserved. We thus explored the gene regulatory network of auditory homeostasis.

Across taxa, ears are delicate mechanoelectrical converters. Their operation can be conceptually divided in a *passive* and an *active* component. The passive system could be approximated by a simple spring-mass oscillator, whose properties are determined by the stiffness of the associated springs or the size of the mass attached to it. In case of the *Drosophila* ear, the chordotonal neurons do not add significantly to the relevant mass of the passive antennal oscillator but they are an important contributor to its passive stiffness ^15^. *K*_*steady*_, the steady-state stiffness of the antennal joint, is a good indicator of this passive stiffness ^13^. Both with regard to its natural life course and the effects of our transgenic manipulations, *K*_*steady*_ remained one of the most stable parameters of auditory function (Fig. 1d and Fig. 4d), suggesting that the causes for the observed functional decline emerge from the active system. The *active* oscillator of the fly’s ear emanates from its auditory transducer modules, i.e. mechanosensory ion channels that act in series with – and receive feedback from ^14^ – probably dynein-based motor proteins ^20^. This functional design explains vast parts of the functional performance of the *Drosophila* ear ^14^; its quantitative modelling also allows for extracting vital parameters of auditory function, such as the amount of energy that auditory neurons inject into the hearing process or the number – and molecular properties – of transducer channels they harbour.

Quantitatively, the hearing loss observed in flies older than 50 days is best described as a loss of the active process, i.e. a loss of power gain (Fig 1c), which injects energy into sound-induced motion of the antennal receiver. This loss of activity is accompanied by a gradual loss of nerve response (CAP amplitudes), which decline steadily from day 25 already, in both males and females (Fig. 1d). Our data shows that ageing occurs on various levels of auditory function, namely transduction, motor-based feedback amplification and signal transformation into action potential responses. Auditory transducers, however, also display a remarkable resilience throughout life; the characteristic nonlinear signatures they introduce into sound receiver mechanics (gating compliances) stay broadly constant up to the age of 70 days (Fig. 1d). First quantitative gating spring model analyses also hint at a possible homeostatic mechanism for this constancy: In both males and females, and across ages, transducer channel numbers were found to be inversely correlated with their respective single channel gating forces. When transducer numbers decrease with age, their single channel gating forces increase, thereby stabilizing the nonlinear mechanics of the sound receiver across the auditory life course (Supplementary Fig. 3). This homeostatic stabilization of receiver nonlinearity is particularly significant, as all changes in receiver mechanics will affect all neurons and thereby global JO function. In order to understand these, and other, homeostatic mechanisms we explored the transcriptional network that mediates them.

We found that 16,243 genes are expressed in the 2^nd^ antennal segment, which harbours the fly’s inner ear (JO); 5,855 out of these change their expression in at least one of the pair-wise age comparisons. Four transcription factors emerged from our bioinformatical analysis as key regulators of the age-variable auditory transcriptome, all of which are conserved in the human genome; these are Onecut, Worniu, Optix and Amos. Given the reported high conversation of binding specificities between fly and human TF orthologues ^38^, these findings are likely to be directly translatable to the human cochlea.

Onecut is a transcription factor known to be involved in photoreceptor differentiation in flies ^39^ and retinal ganglion cell development in mice, where it cooperates with Pou4f2 (*acj6*) and Atoh7 (closest fly orthologues: *ato, amos*) ^40^. We here report an essential role for Onecut in fly hearing or - more precisely – in the homeostatic maintenance of fly hearing. An adult-specific knockdown (KD) of the *onecut* gene across JO neurons affects all levels of auditory system function and leads to deafness. The *onecut* KD phenotype includes near complete losses of auditory (i) transducer gating, (ii) amplification and (iii) nerve responses, as well as (iv) a loss of sound-evoked behaviour. A first probing of possible Onecut targets through qPCR in *onecut* KD flies (Fig. 4f) might reveal one possible mechanism of action, which is the direct transcriptional regulation of the two interdependent TRPV channels Nan and Iav. Nan/Iav are thought to form a heterodimeric ion channel specifically in chordotonal neurons. Both genetic ^35,36^ and pharmacological ^41^ ablations of Nan/Iav channels have been shown to eliminate chordotonal mechanosensory function. After an adult-specific knockdown of *onecut*, JO expression levels of both *nan* and *iav* showed a dramatic decline. This downregulation coincided with a near complete abolition of the gating compliances associated with sound-sensitive neurons, indicating a failure of auditory transduction. A total loss of transducer function would also be sufficient to explain the effects on auditory amplification and nerve responses observed further downstream the auditory signalling chain. Interestingly, both Nan/Iav ^35,36,42^, as well as NompC ^43,44^, have previously been proposed as auditory transducer components in *Drosophila*. Elegant further studies also demonstrated beyond doubt that NompC contains all elements required to form a bona-fide mechanotransducer channel ^45,46^. In contrast to *nan*/*iav*, however, *nompC* transcript levels in JO remained unchanged after *onecut* KD.

*worniu* (*wor*) is a zinc finger transcription factor that belongs to the Snail family. We here demonstrate that adult-specific down-regulation of *wor* enhances auditory amplification and sharpens auditory tuning. These effects are sustained up until 30 days of downregulation (at 30°C) - when the ears of control flies already show a near complete loss of power gain - suggesting that knockdown of *worniu* can protect distinct aspects of auditory function from their age-dependent decline. Genes previously reported to be upregulated in *wor* mutants included cadherins and trafficking proteins, e.g. Rabs ^47^; our bioinformatics prediction also support a role of wor in the regulation of the cellular trafficking machinery (Supplementary Fig. 2). In JO, the adult-specific KD of *wor* had virtually no effect on auditory transducer gating or the transformation of antennal motion into nerve responses. Auditory amplification and tuning sharpness, however, were significantly enhanced in *wor* KD flies; both of these parameters are linked to the dynein-based motor machinery that acts in series with the auditory transducer channels. Consistent with these mechanistic considerations, qPCR analyses of the JOs of *wor* KD flies showed a downregulation of Dhc98D; neither *nompC*, *nan* nor *iav* levels were affected.

Optix belongs to the sine oculis homeobox (SIX) family of transcription factors and is required for eye formation ^48^. The adult-specific knockdown of *Optix* in JO neurons leads to an increase in the receiver’s power gain and a shift of its best frequencies to lower values, both indicative of more active system. qPCR analyses showed that these changes in auditory activity coincided with an upregulation of *nompC*. These relations are consistent with the reported roles of NompC in auditory amplification ^37^. Auditory transducer gating, however, was not affected; both numbers and single channel gating forces of sensitive auditory transducers were identical between the ears of Optix KDs and control flies. Also, CAP responses to small antennal displacements (as caused by auditory stimuli) were unchanged. CAP responses to larger displacements (as caused by non-auditory stimuli), in contrast, were decreased as compared to controls. This suggests a more complex role of Optix in the homeostatic regulation of auditory, as well as non-auditory populations of JO neurons.

*amos* is a proneural gene from the family of basic-helix-loop-helix (bHLH) transcription factors. bHLH transcription factors also include *ato*, which specifies chordotonal organs ^18,49^, R8 photoreceptor precursors ^49,50^, and a subset of olfactory sense organs ^51^. *amos* specifies two other subsets of olfactory sense organs and a mechanosensory subset of larval bipolar dendritic neurons ^52,53^. *Ato* and *amos* share a high sequence similarity in their bHLH domains (73% amino acid identity) and their basic - DNA-binding - regions are identical. Maung and Jarman showed that *amos* is capable to rescue eye development independent of *ato* ^54^. Weinberger et al. showed that the coding sequence of *amos*, when used to replace the coding sequence of *ato*, is sufficient to produce a fully functional *Drosophila* ear, the performance of which is statistically identical to the native, *ato*-induced organ with respect to all quantitative parameters also used in this study ^55^. While *amos* was found to be expressed in adult JOs, no such expression was found for *ato*. Consistent with this finding, an adult-specific knock-down (KD) of *ato* does not have any significant effect on fly hearing (Supplementary Table 6). The KD of *amos*, in contrast, produces an accelerated hearing loss phenotype that is characterised by a loss of power gain and tuning sharpness. Interestingly though, best frequencies of *amos* KD receivers do not move towards the passive system but instead show a small - but significant - move in the opposite direction, indicating a larger independence between fundamental parameters of *Drosophila* hearing than appreciated by current models ^14,56^. *amos* KD also leads to a loss of nerve responses in high-threshold units of JO and a homeostatic reorganization of sensitive transducer channels, characterised by a slight decrease in single channel gating forces and a slight increase in channel number (Fig. 4d). Consistent with bioinformatical predictions, qPCR analyses of JOs of *amos* KD flies showed significantly reduced expression levels for the here newly described auditory dynein Dhc98D but no effects on the three auditory TRP channels tested (nompC, nan, iav).

Interestingly, all four master regulators are predicted to act upstream of phototransduction genes (Fig. 2b), including visual opsins, which have been previously linked to *Drosophila* auditory function ^21^ and ciliary maintenance ^57^ and are also upregulated during auditory ageing (Fig. 2a).

Our study has identified novel master regulators of auditory maintenance, some of which work as bidirectional homeostatic actuators within the fly’s auditory neurons. If the regulator’s upregulation, e.g., results in an improvement of a specific auditory function, then its downregulation leads to a worsening (*amos*), or vice versa (*Optix*). Future experiments will shed more light on the downstream targets of these master regulators and their specific roles in hearing and auditory homeostasis. Some key conclusions, however, can be drawn already. All four homeostatic master regulators that emerged from our screen are evolutionary conserved; they either form key regulators of specific sensory (or neural) tissues or constitute paralogs of such regulators; their predicted (and in part validated) regulons, however, do not comprise of classic developmental genes (such as proliferation, apoptosis) but rather of known effector genes for specific auditory functions, e.g. ion channels and motor proteins. This suggests a scenario where developmental and homeostatic functions are divided between pairs (or groups) of paralogs. Examples for such pairs from our study would be *ato*/*amos* or *ct*/*onecut*. Sometimes, as is the case for the proneural master gene *ato*, the homeostatic roles seem to have been fully transferred to a paralog (*amos*). If true beyond *Drosophila*, this could also have implications for the design of gene-therapeutic trials to reverse human hearing loss, which currently concentrate on key developmental genes - such as e.g. ATOH1. ATOH1’s ‘next of kin’ – such as e.g. ATOH7 or NEUROD1 – might be worth having a look at.

## Materials and Methods

### Fly stocks and husbandry

To assess the natural life course of hearing in *Drosophila* the following lines were used as wildtype references: Canton-S line (Bloomington), Canton-S (Goodwin lab), Canton-S (Kamikouchi lab), Oregon-R.

To probe the expression of predicted transcription factors the following lines used: Fly-TransgeneOme (fTRG) sGFP tagged lines from VDRC^58^ for *amos*, *onecut* and *Optix* (*optix:GFP 318371/10042*), *wor*-Gal4 (kindly provided by A. Carmena).

*elav*-Gal4; UAS-RFP-nls/+; Mi{PT-GFSTF.0}alphaTub85E[MI08426-GFSTF.0]/+ was used to monitor JO neurons across the flies’ lifespan.

y[1]w[*]; tub-Gal80ts; NP0761 was used for adult-specific downregulation (via RNAi knock-down) or upregulation (via overexpression) of target genes across all JO neurons.

RNAi lines were obtained from the Bloomington *Drosophila* Stock Centre (BDSC) and Vienna *Drosophila* Research Centre (VDRC). Attp2 and attp40 served as control lines for the TRIP collection and VDRC 6000 was used as control for the KK lines.

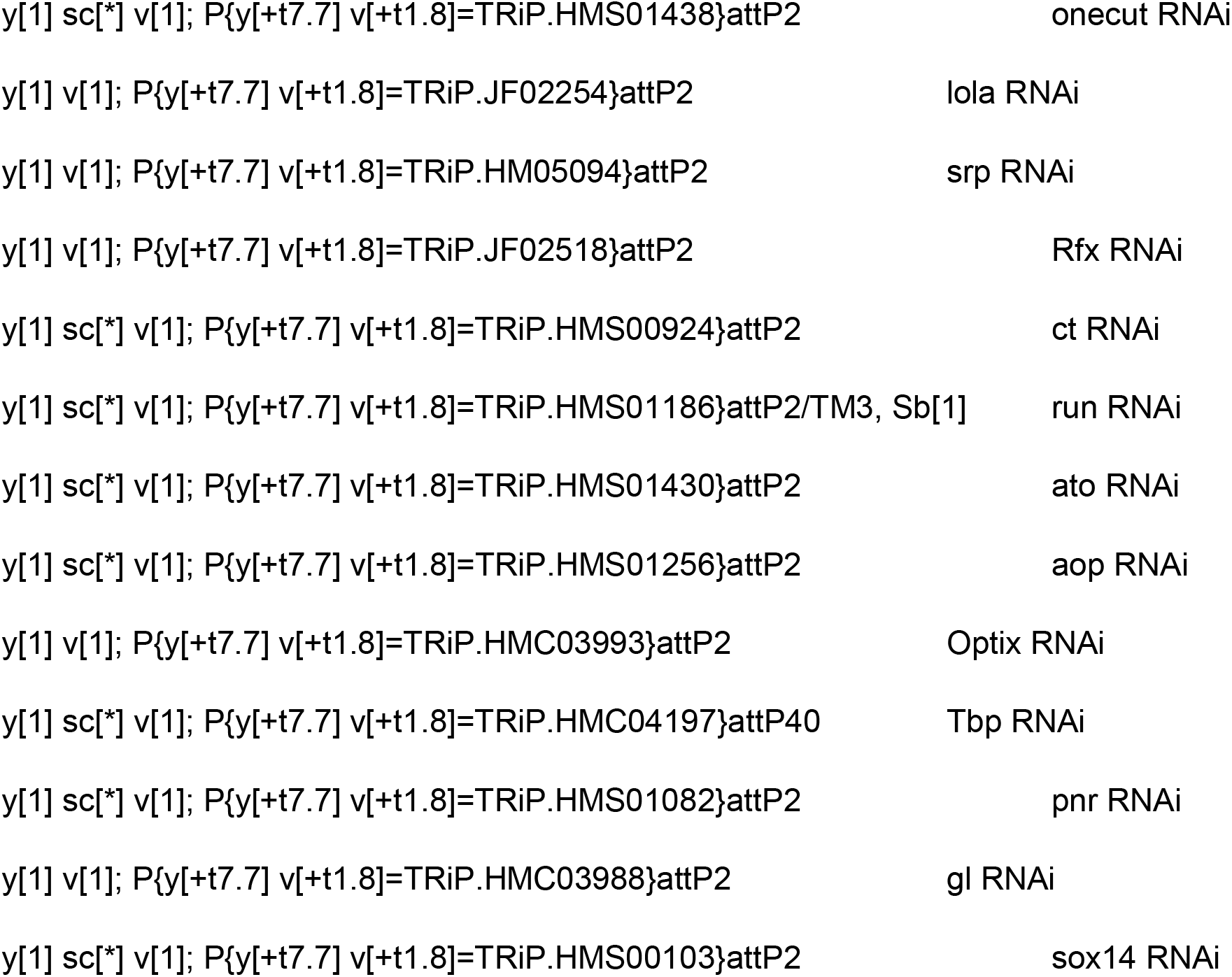
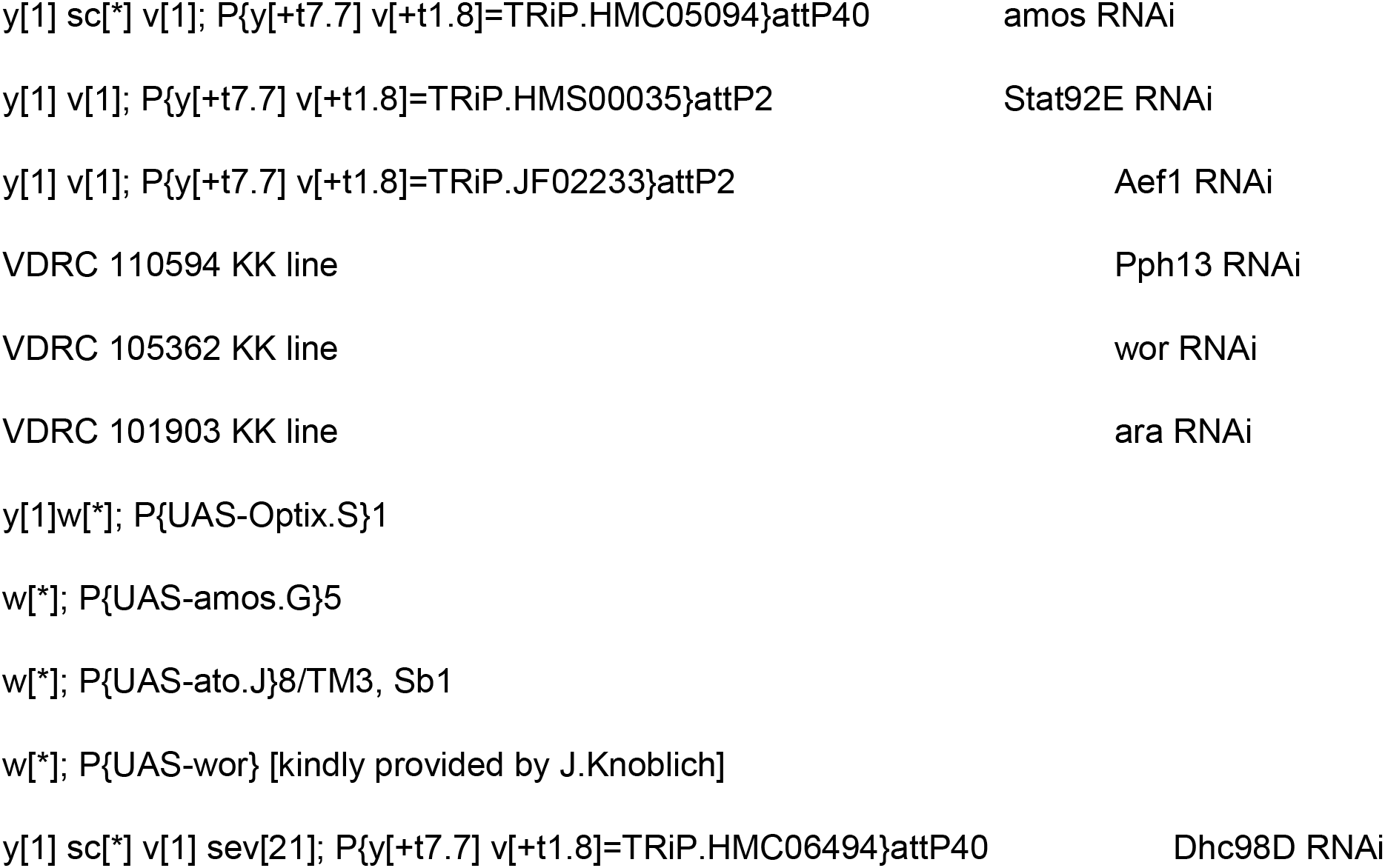

### Immunostainings of JO

Fixation and immunostainings followed standard procedures. Briefly, 10 days old adult female heads were dissected in PBS, fixed with a 4% formaldehyde solution (in PBT) for one hour while rotating at room temperature (RT); three heads were placed exposing the antennae into silicon blocks filled previously with hot gelatin-albumin mixture. Silicone blocks were then quickly cooled down at 4°C for 10 minutes and incubated with 6% formaldehyde solution overnight at 4°C. Thereafter, silicone blocks were extracted and incubated further with Methanol for 10 min at RT, before being washed with PBS for 30 min at RT. 30 µm vibratome sections were cut using a vibratome (Ci 5100mz, Campden Instruments) and antennae sections collected in PBT (PBS with 0.3% Triton X-100) and afterwards washed three times for 15 min at RT. After blocking for 1 hr at RT (blocking solution: PBS with 1% Triton X-100, 2% BSA, 5% normal goat serum), samples were incubated with primary antibodies in blocking solution overnight at 4°C, then washed again three times in PBT and incubated with secondary antibodies diluted in blocking solution for 2 hr at RT. Samples were then washed again three times in PBT and, finally, briefly washed in PBS before mounting. Primary antibodies used in this study are:

Rb anti-GFP 1:1000 (Life Technologies), Rat anti-elav 1:250 (Hybridoma Bank), Goat anti-HRP∷Cy3 1:500 (Jackson ImmunoResearch). Secondary antibodies conjugated with Alexa 488, and Alexa 633 (Life Technologies) were used at 1:500. All samples were mounted in Dabco (Molecular Probes, H-1200). Images were acquired with a LSM 510 Zeiss confocal microscope with a Plan-Neofluar 40x/1.3 Oil objective. Z-stacks (optical slice thickness: 1μm) were taken to image throughout Johnston’s organ (JO). Images were assembled and analysed in ImageJ (Fiji).

### Neuron counts

Flies of genotype *elav*-Gal4; UAS-RFP-nls/+; Mi{PT-GFSTF.0}alphaTub85E[MI08426-GFSTF.0]/+ were aged at 25°C. Fly antennae of day1, day5, day25 and day 50 flies were dissected in PBS, such that left and right antennae remained attached to the cuticle, and that the third antennal segment and the associated arista remained intact. Antennae were then briefly fixed for 10 minutes in 4% formaldehyde in PBS, washed three times in PBT and finally mounted in glycerol. Fly JO-s were imaged with an LSM 510 Zeiss confocal microscope with a Plan-Neofluar 40x/1.3 Oil objective. Z-stacks (optical slice thickness: 1μm, 80 slices in total) were taken to image throughout Johnston’s organ (JO). Images were processed and unspecific background removed using the FluoRender programme. Single Z-stacks were processed in ImageJ. The Eve programme (with kind permission from Kei Ito)^23^ was used to count neurons automatically (XY:Z ratio was set depending on the number of the stacks), neuron radius was set to 4 and Bending 1 at 300 cells was used as a cut-off. Processed images were saved as new files (including cell count information) and result files were produced. Number of cells was corrected manually by using the ImageJ cell counter plugin.

### RNA sequencing

Male and female Canton-S flies of different ages (days 1, 5, 10, 25, and 50) were anesthetised on ice, their second antennal segments dissected and collected in Lysis Buffer (containing 1% β-mercaptoethanol, as provided in the Qiagen RNeasy Mini Kit). As soon as dissections were completed for a given time point, samples were frozen at −80°C. When enough samples were collected, RNA was extracted according to the Qiagen RNeasy Mini Kit protocol.

Reverse transcription and pre-amplification were carried out with the SMART-Seq v4 Ultra Low Input RNA Kit for Sequencing (Clontech). All samples were quality controlled and cDNA concentrations measured with an Agilent BioAnalyzer 2100. Sample libraries were prepared with a Nextera XT DNA Library Preparation kit (Illumina). Thereafter, paired-end 75bp reads were sequenced on an Illumina NextSeq 500 platform.

The RNAseq .fastq files were aligned in the Partek Flow software to the most recent version of the *Drosophila* genome (dm6) obtained from the Berkeley *Drosophila* Genome Project at UCSC.

In order to generate raw sequence counts, .bam files created in Partek software were processed in HTSeq. These counts were then used in DESeq (105) in R/Bioconductor to measure differential expression across genes and for conducting ANOVA statistical analyses of each comparison. Further data filtering took place to reduce the maximum false discovery rate (FDR) to 10% limiting the expression fold change threshold to ±1.5x.

### Quantitative PCR (qPCR)

Flies of different genotypes were collected and frozen immediately with liquid nitrogen and then kept at −80°C. After 50 flies were collected they were vortexed and second antennal segments were collected (100 antennae) in Lysis Buffer containing 1% β-mercaptoethanol (as part of the Qiagen RNeasy Mini Kit). In accordance with the Qiagen RNeasy Mini Kit protocol, RNA was extracted immediately and RNA samples were then kept at −80°C. Reverse transcription was carried out with the High Capacity RNA to cDNA kit (Applied Biosystems) following the manufacturer’s protocol.

In order to proceed to pre-amplification with TaqMan PreAmp Master Mix Kit, a “pooled assay” of Taqman primers was prepared (containing probes for the target genes of interest, i.e. *nompC*, *nan*, *iav*, *Dhc98D*). TaqMan probes of interest were mixed together and diluted 1:100 in TE buffer. The pre-amplification procedure followed the manufacturer’s protocol. The pre-amplified cDNA was diluted 1:20 in RNAse and DNAse-free water and the qPCR was performed in the immediate aftermath. qPCR assays were run on a Step One Plus ABI machine. Prior to the reaction, the 96 well plate set up was designed with help of the Step One Plus software. Three negative controls were run per target as well as three replicates for each sample and each target. *SdhA* was chosen as endogenous control as one of the housekeeping genes that has the most stable expression at different ages ^59^.Reactions were prepared according to the TaqMan Gene Expression Assay protocol.

Cycle threshold (Ct) values were extracted from the Step One Plus Software data analysis. The ΔΔCt and relative quantification were calculated in Excel as follows:

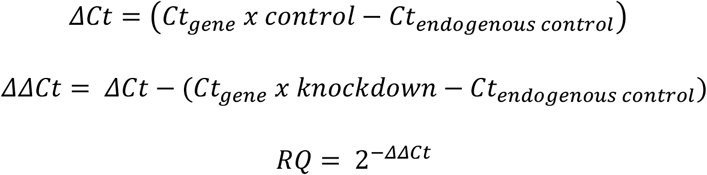

The three RQ values were averaged for each biological replicate and standard deviations generated in Excel. At least three biological replicates were performed for each experiment. Statistical were performed in SigmaPlot.

### Transcription factor prediction – bioinformatical analysis in iRegulon

The iRegulon plug-in ^32^ was used in the Cytoscape software to predict transcription factors/regulators based on their binding motifs. A list of genes of interest was submitted and predicted transcription factors were then selected according to their normalised enrichment scores (NES) for a particular motif, or group of motifs, within the list originally submitted to iRegulon.

### Gene ontology analysis - GORILLA

The online interface GOrilla (**G**ene **O**ntology en**RI**chment ana**L**ysis and visua**L**iz**A**tion tool) was used to classify genes of interest according to their gene ontology ^28,29^

Functional classifications were generated for biological processes and molecular functions. Enrichment scores were calculated:

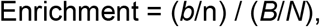

where *b* is the number of genes in the intersection, *n* the number of genes in the target set, *B* the total number of genes associated with a specific GO term and *N* the total number of genes.

### Analyses of JO function

Flies were raised on standard medium in incubators maintained at 25°C and 60% relative humidity (RH), with a 12 hr:12 hr light:dark cycle. Three Canton-S lines, and one Oregon-R line were used for the ageing experiments. Female and male flies were collected on the day of eclosion using CO_2_ sedation and allowed to age in separate vials at 25°C for 1, 5, 10, 25, 50, 60 and 70 days – free fluctuation assays and electrophysiology experiments were conducted at room temperature (21°C −22°C). Adult-specific RNAi knock-down mutants (whose larvae and pupae were kept at 18°C in order to repress the Gal4-mediated transcriptional activation via a Gal80ts repressor) were collected on the day of eclosion and transferred to 30°C (for maximal activation of the Gal4/UAS expression system), 60% RH and kept at 12 hr:12 hr light:dark cycles for 2 weeks prior to the experiments. Flies were mounted as described previously ^13^. Briefly, flies were attached, ventrum-down, to the head of a Teflon rod using blue light-cured dental glue. The second segment of the antenna under investigation was glued down to prevent non-auditory background movements. The antenna not under investigation was glued down, completely abolishing any sound-induced motion and interference with the contralateral recordings. An active vibration isolation table (model 63-564; TMC, USA) was used. After mounting, flies were oriented such that the antennal arista was perpendicular to the beam of a laser Vibrometer (PSV-400; Polytec, Germany) and free fluctuation recordings could be taken. Electrostatic actuation used two external actuators positioned close to the arista (for details see Effertz et al, 2012 ^43^). Two electrodes were inserted into the fly – a charging electrode was placed into the thorax so that the animal’s electrostatic potential could be raised to −20 V against ground, and a recording electrode for measuring mechanically evoked compound action potentials (CAPs) was introduced close to the base of the antenna under investigation. The charging electrode also served as reference electrode for the CAP recordings.

Arista displacements were measured at the arista’s tip using a PSV-400 LDV with an OFV-70 close up unit (70 mm focal length) and a DD-500 displacement decoder. The displacement output was digitized at a rate of 100 kHz using a CED Power 1401 mk II A/D converter and loaded into the Spike 2 software (both Cambridge Electronic Design Ltd., Cambridge, England). Free (i.e. unstimulated) fluctuations of the arista were recorded both before and after the experiment to monitor the physiological integrity of the antennal ear. Only those flies, which maintained a stable antennal function throughout the experiment (maximally allowed change of best frequencies: 20%) were analyzed.

### Tests of sound-evoked behaviour

*Drosophila melanogaster* males increase locomotor activity in response to a playback of courtship songs ^24^. We exploited this phenomenon to test hearing across the *Drosophila* life course. To conduct measurements, flies were housed in 5×65mm Pyrex glass tubes. One end of the tube was sealed with an acoustically transparent mesh, the other end contained food consisting of 5% sucrose and 2% agar medium covering ~ ¼ of the tube. Glass tubes were then loaded into high-resolution *Drosophila* activity monitors (MB5; Trikinetics, Walham, USA). MB5 monitors harbour 17 independent infrared (IR) beams bisecting each tube at 3mm intervals, allowing for a high-fidelity recording of the flies’ activity. Detectors were set to count all beam breaks occurring within one minute for the duration of each experiment. Activity counts were registered independently at each beam position within a tube; all beam breaks, irrespective of beam position, were then pooled. This procedure allowed for calculating the total activity of all flies in that tube. To maximise data collection, three MB5 monitors were stacked together forming a grid allowing to record from 36 tubes (totalling 108 flies) simultaneously over the course of a single experiment. The MB5 activity monitors were placed centrally in front of a 381mm wide bass speaker (Eminence Delta 15, 400W, 8 ohm) with the tubes’ acoustically transparent mesh facing the speaker at a distance of ~60mm from the speaker membrane. The speaker was connected to an amplifier (Prosound 1600W). To avoid interference from non-air-borne vibrations, the MB5 monitors – but not the speaker - were placed on a vibration isolation table. Sound stimuli were adjusted to reach peak amplitudes of 90 dB SPL at the middle of the monitor tubes. Courtship stimuli played at these intensities are known to elicit reproducible behavioural responses in males ^60^. The sound stimulus (played, and controlled from the Spike2 software) consisted of a ‘master pulse’ that was repeated to form 2s long trains with 40ms interpulse intervals (IPIs). The master pulse was generated by averaging previously recorded original courtship song pulses (~1000 pulses from 10 Drosophila melanogaster males). Each pulse train was followed by a 2s long silence; this elementary kernel was played continuously for 15 minutes at the beginning of every hour. The 15 min of sound stimulation were played in loop with 45 min of silence for an entire circadian day (24h at a 12-hour light, 12-hour dark cycle). Responses for each hour were then collapsed into a single median response to cancel out any circadian variations of responsiveness.

Activity displayed during *stimulus presentation* was determined by the sum of all activity displayed during the first 15 minutes of every hour (i.e. during the phase of sound stimulation) and averaged over the whole experimental day. *Baseline activity* was determined by the sum of all activity displayed during the last 15 minutes of every hour (i.e. during the silent phase directly preceding the next stimulus phase) and also averaged over the whole experimental day.

The room, in which the recordings took place was held at a constant 25°C temperature (@~40% RH) and followed a 12-hour light, 12-hour dark cycle, which was kept consistent with the flies’ entrainment regime prior to experiment start. Exposure to courtship sound is known to induce male flies to also produce mating songs. To prevent these stimulus-induced mating songs from interfering with the sound stimulus, experimental flies were anaesthetized using CO_2_ and their wings clipped 2-4 days prior to the initiation of the experiment. At least 2/3 of the wing was removed using microdissection scissors. After allowing time for healing post procedure, the flies were again CO_2_-anesthetized and transferred into the glass tubes before being placed into the MB5 monitors. For each experiment, flies were exposed to the sound stimulus as soon as they were placed into the monitors; however, only data recorded from the first light transition on was used for analysis. This allowed flies to have ~12hr to adapt to the stimulus, the new environment and to recover from after-effects of CO2 exposure. After this equilibration stage, data was collected for 48 hours.

**Table 1. Previously identified JO genes with age-variable expression** The highlighted area (orange) shows genes that are changing its expression mainly in males, ‘Avg exp’ means ‘Average expression’. 36.7 % (108 out of 294) of all previously reported JO genes show age variable expression patterns. Please note that many genes previously identified ^21^, such as rhodopsins, the mechanosensitive ion channel Nan, the ATP pumps nervanas, innexins, tilB etc., show high variability in JO across ages.

**Table 2. Mouse genes linked to deafness, which are conserved - and expressed - in the *Drosophila* JO**. 65% (65 out of 100) of all reported mammalian/human hearing loss genes are conserved in *Drosophila* and expressed in JO. Genes that change their expression in one of the two age comparisons in both sexes are shown in bold type. Novel candidate genes for mammalian/human hearing loss, recently identified in a large scale hearing loss screen ^10^ are highlighted in light orange, others identified by Ingham and colleagues ^11^ are highlighted in blue, and Age-related deafness genes identified by Potter et al. ^12^ are highlighted in green. ‘Avg exp’ stands for ‘Average expression’.

## Supporting information

Suppl figures adn tables

