## Supplementary material for "Homeostatic maintenance and age-related functional decline in the *Drosophila* ear": Suppl figures adn tables

#### Supplementary legends (Figures and Tables)

**Supplementary Figure 1. JO neuronal count across the life course.** (a) The number of JO neurons does not change between days 1, 5, 25 and 50. From day 1 to day 50 – i.e. during the time of homeostatic equilibrium – neuronal numbers stay constant. Dotted red line indicates previously published number of neurons (477±24; [23]), dashed lines indicate standard deviations. (b) Whole-mount preparation of JO expressing the neuronal marker Elav (magenta) and support cell marker αTub85E (green). Whole-mount imaging of JO was used to count JO neurons at different ages.

**Supplementary Figure 2. Prediction of the downstream regulon using *i*Regulon.**

(a)Trafficking genes are predicted downstream of Wor. (b) Receptors (and ion channels), including NompC and Nan (shown in yellow) are predicted downstream of several transcription factors, including onecut, Optix, Ci, Sox110B, Pph13, ato and Dr. (c) Structural genes are predicted to be largely downstream of *amos* and *wor*. (d) Motor proteins belonging to the dynein family, are predicted downstream of *amos* and wor; this also includes dyneins previously identified in JO (shown in yellow) [20,21].

**Supplementary Figure 3. Homeostatic link between numbers and gating forces of transducer channels preserves sound receiver nonlinearity.** (a) For both the sensitive (left, blue symbols) and the insensitive (right, green symbols) population of transducers, the relation between channel number (N) and single channel gating force (z) follows a simple model (dashed line), which assumes a constant contribution of the respective transducer population to the overall receiver nonlinearity (corresponding to $N \sim z^{-2}$, see also [13]). [Summary data, pooled from all ages and both sexes]. (b) Shown are the numbers (green) and gating forces (red) for sensitive (upper) and insensitive (lower) transducer channels of the individual ages tested, for both females (left) and males (right). Note the age-related changes in both channel numbers and gating forces and the constancy of their resulting contribution to the sound receiver's extent of nonlinearity (yellow).

**Supplementary Figure 4.** Over-expressions of *Optix* for 15 days and *amos* for 30 days revert their respective knock-down phenotypes (compare to the Figure 4a).

**Supplementary Table 1. *Drosophila* auditory mechanics across the life course.**

Three principal parameters of sound receiver function were assessed in Canton-S and Oregon-R flies at different ages (days 1, 5, 10, 25, 50, 60 and 70): (i) $f_0$ , the receivers' best frequency [in Hz]; (ii) the receivers' tuning sharpness or 'quality factor' *Q* [dimensionless] and (iii) the receivers' energy gain [in $k_BT$]. Pair-wise student t-tests or Mann-Whitney (MWUS) tests were used to assess statistical significance (choice of test depending on data distribution). Significant changes are highlighted in yellow. Canton-S (K) and Canton-S (G) are two other Canton-S lines kindly provided to us by Azusa Kamikouchi and Stephen Goodwin, respectively. Both males and females were measured, except for Oregon-R, where only females were measured.

**Supplementary Table 2. JO RNA-Seq transcriptomic data across the lifespan for both males and females.**

Flybase ID numbers, gene annotated names, gene-ontological description and the genes' chromosomal positon are shown for all genes detected in the RNAseq data. Average normalised expression counts are shown in the “baseMean” column, log2Fold changes (log2FC) are shown in pair-wise comparisons between ages of day 50 to day 1 (“log2FC_d50m_d1m”), day 25 and day 1 (“log2FC_d25m_d1m”), etc. *p*-adjusted values are shown in the column “padj”. The analysis was done by using ANOVA. Data is shown for both males (“m”) and females (“f”). Raw counts are shown for each gene at the end of the table, where m1A stands for day1 male replicate number 1, f5B – for female day 5 replicate 2.

**Supplementary Table 3.** Age-variable transcriptome of JO

Three comparisons were performed of the day 1 to day 5, day 5 to day25 and day 25 to day 50 for genes that were changing in both males and females. The analysis was done by using ANOVA.

Flybase ID numbers, gene annotated names, gene-ontological description, mouse homologues and the gene positon in the chromosomes are shown for all genes of the JO RNAseq data set. Average normalised expression counts are shown in the “baseMean” column. Inclusion criteria were: Log2Fold-Change at least 1.5 times with False Discovery Rate (FDR) of 10%, at least in one of the three comparisons. Signs indicate whether a respective gene was downregulated (negative) or upregulated (positive) for the respective age pair.

**Supplementary Table 4. Summary list of age-variable JO genes identified through RNA-Seq-based transcriptomics.**

JO genes that are either up- or down-regulated in between two age pairs. Age pair 1: (i) day 5 and day 25; age pair 2: (ii) day 25 and day 50. JO genes with significant expression changes are highlighted in red (up-regulated) or blue (down-regulated). Example: genes that are upregulated from day 1 to day 5 with a log2fold change between 2 and 4 are shown as “day 1to5 log2fold 2-4” and highlighted in red. Genes that are down-regulated from day 25 to day 50 with a log2fold change between -1 and -2 are shown as “day25to50 log2fold -2 to-1” and highlighted in blue.

**Supplementary Table 5. 37 iRegulon predicted transcription factors (TFs) expressed in the *Drosophila* JO.** Genes that change their expression in one of the two age comparisons in both sexes are shown in bold type. The 19 genes functionally probed in our study are highlighted in light orange. ‘Avg exp’ stands for ‘Average Expression’.

**Supplementary Table 6. Auditory mechanics after overexpression (OE) or knockdown (KD) of transcriptional regulators.**

Adult-specific manipulation of transcriptional regulators was conducted by using the temperature sensitive repressor tubulin-Gal80[ts] together with the NP0761 Gal4 driver, which drives expression in all JO neurons. Knock-down (KD) or overexpression (OE) of regulators was achieved by using UAS-RNAi or UAS cDNA constructs, respectively. KD or OE constructs were driven for 15 and/or 30 days at $30^{0}$C and then hearing was assessed through free fluctuation analysis of unstimulated sound receivers as described previously [43]. Three principal parameters of sound receiver function were assessed: (i) f0, the receivers’ best frequency [in Hz]; (ii) the receivers’ tuning sharpness or ‘quality factor’ Q [dimensionless] and (iii) the receivers’ energy gain [in kBT]. Pair-wise student t-tests or Mann-Whitney (MWUS) tests were used to assess statistical significance (choice of test depending on data distribution). Both males and females were measured.

### Supplementary Figure 1.

a

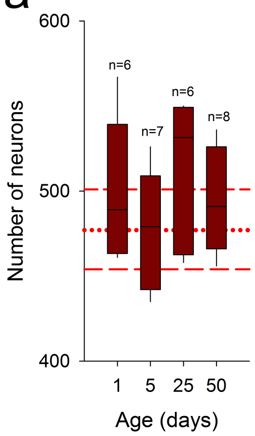


b

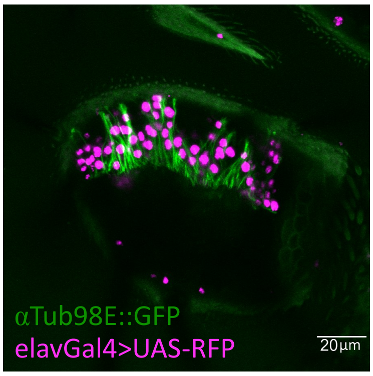

Supplementary Figure 2.

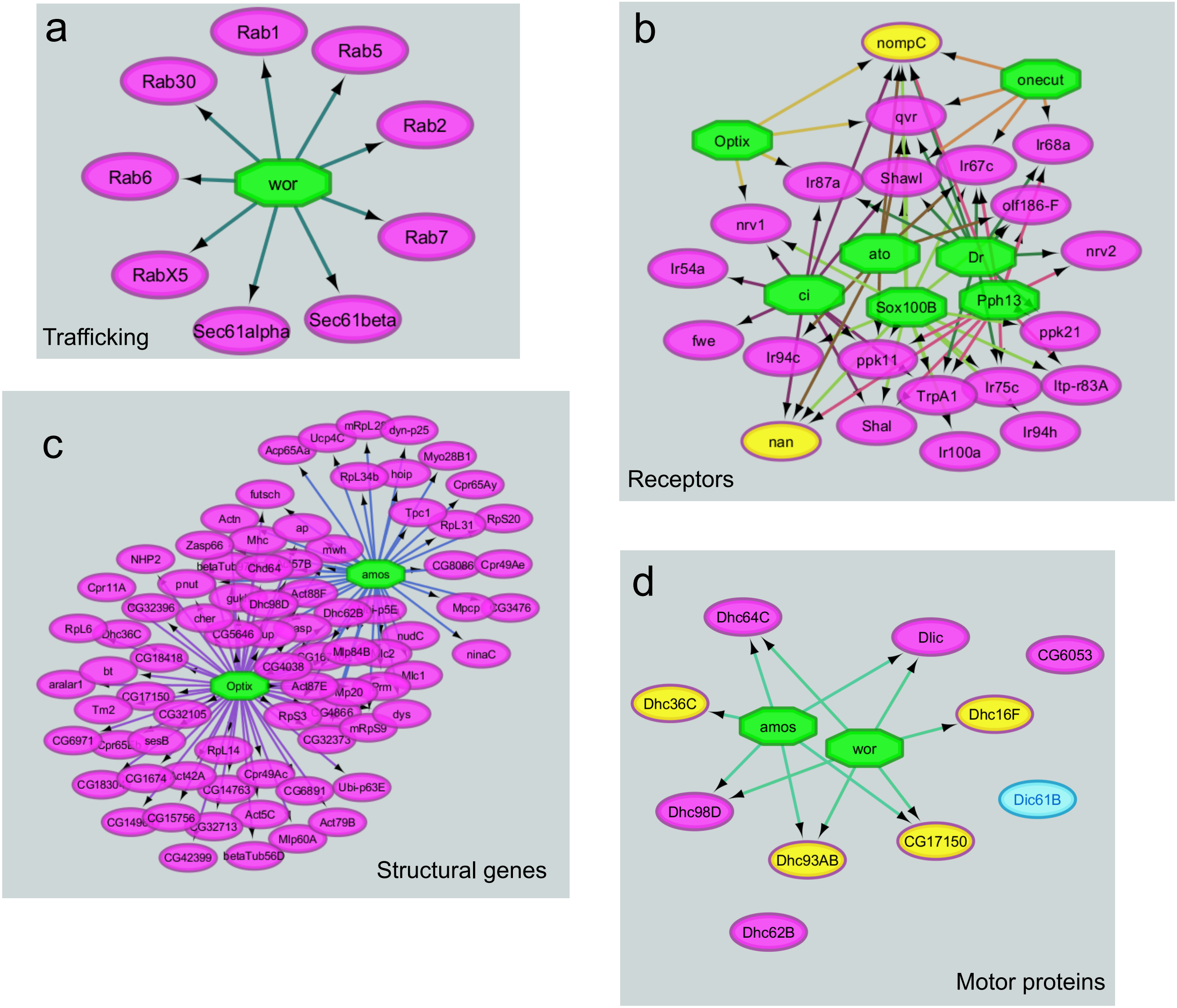

### Supplementary Figure 3.

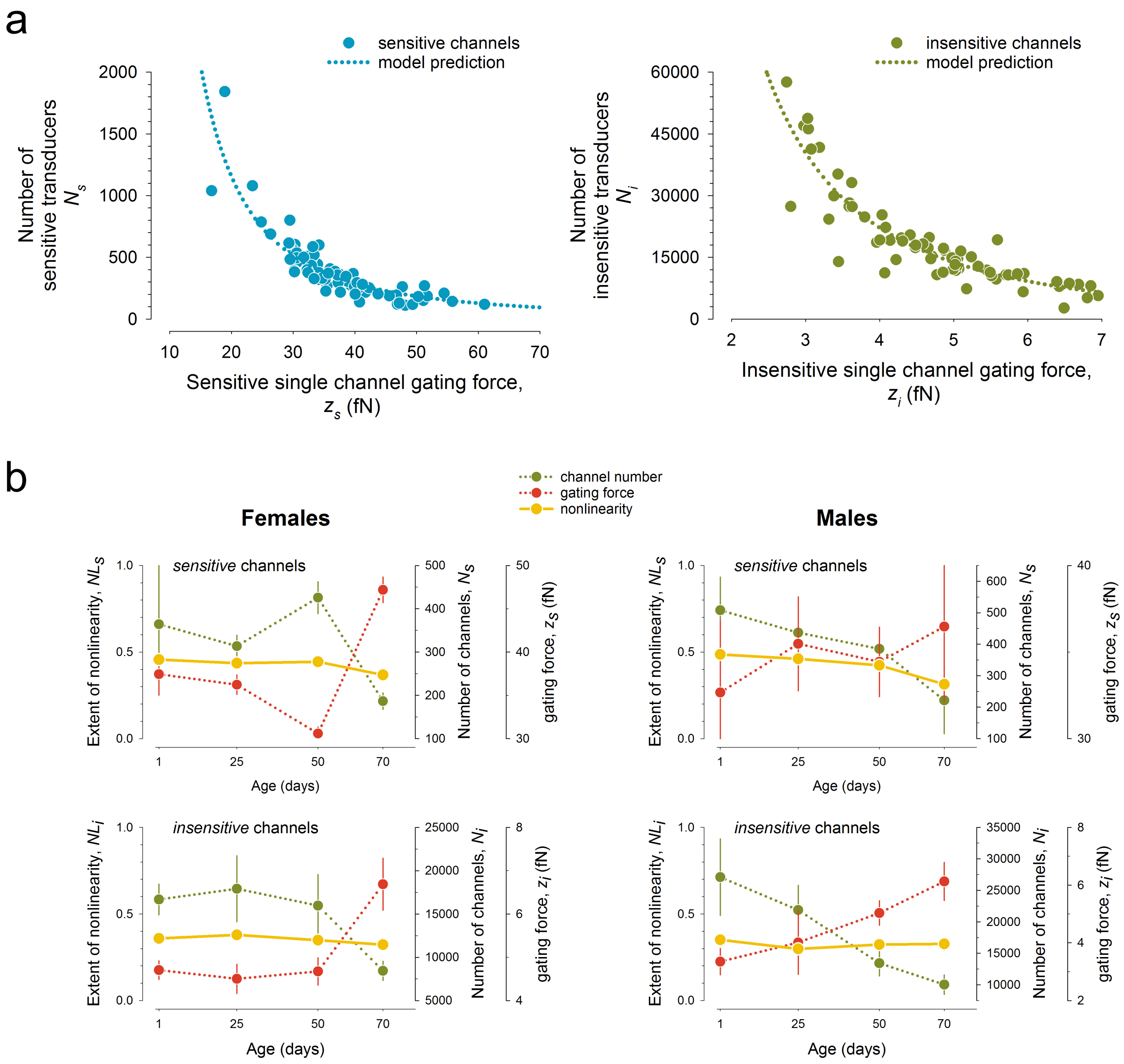

### Supplementary Figure 4.

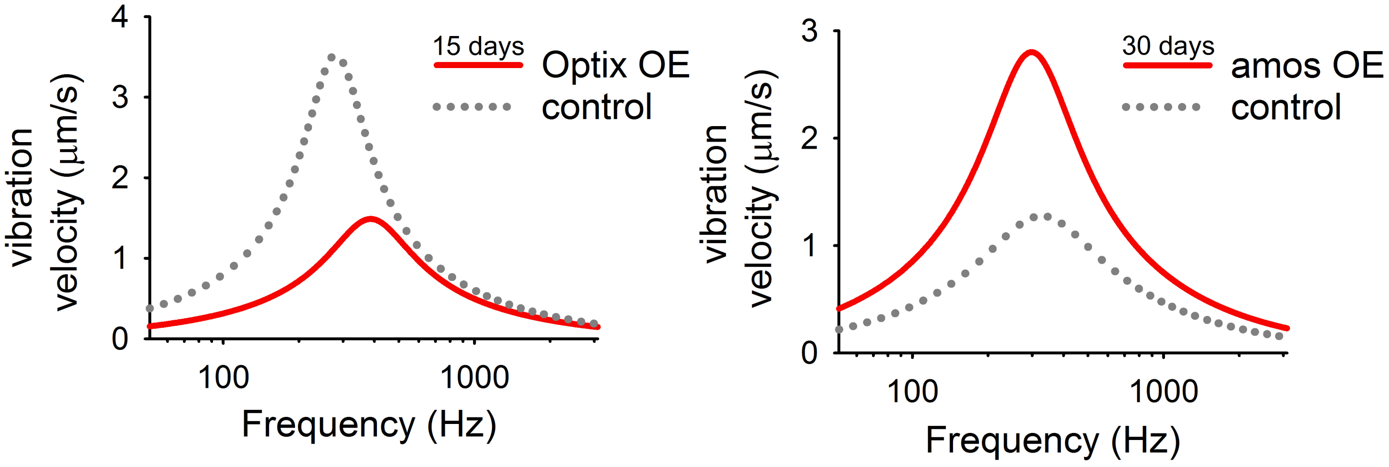

**Supplementary Table 1.** ***Drosophila*** **auditory mechanics across the life course.**

Three principal parameters of sound receiver function were assessed in Canton-S and Oregon-R flies at different ages (days 1, 5, 10, 25, 50, 60 and 70): (i) f0 , the receivers' best frequency [in Hz]; (ii) the receivers' tuning sharpness or 'quality factor' Q [dimensionless] and (iii) the receivers' energy gain [in kBT]. Pair-wise student t-tests or Mann-Whitney (MWUS) tests were used to assess statistical significance (choice of test depending on data distribution). Significant changes are highlighted in yellow. Canton-S (K) and Canton-S (G) are two other Canton-S lines kindly provided to us by Azusa Kamikouchi and Stephen Goodwin, respectively. Both males and females were measured, except for Oregon-R, where only females were measured.

| Genotype | age | sex | Parameter | N | Mean | Median | StDev | StErr | p-value |
|---|---|---|---|---|---|---|---|---|---|
| CantonS | day1 | male | f0 [Hz] | 18 | 229.63 | 217.41 | 74.29 | 17.51 | -- |
| | | | Q | | 2.81 | 2.21 | 2.04 | 0.48 | -- |
| | | | Energy [kBT] | | 23.15 | 20.17 | 8.89 | 2.09 | -- |
| | | female | f0 [Hz] | 17 | 197.26 | 190.25 | 45.31 | 10.99 | -- |
| | | | Q | | 2.30 | 2.00 | 0.99 | 0.24 | -- |
| | | | Energy [kBT] | | 20.69 | 18.45 | 8.31 | 2.02 | -- |
| | day5 | male | f0 [Hz] | 13 | 262.61 | 264.72 | 56.09 | 15.56 | 0.032 (MWUS) |
| | | | Q | | 3.03 | 2.94 | 1.33 | 0.37 | 0.389 |
| | | | Energy [kBT] | | 17.27 | 16.37 | 8.28 | 2.30 | 0.048 (MWUS) |
| | | female | f0 [Hz] | 8 | 238.43 | 232.90 | 29.47 | 10.42 | 0.021 (MWUS) |
| | | | Q | | 2.62 | 2.37 | 1.07 | 0.38 | 0.432 |
| | | | Energy [kBT] | | 14.54 | 13.07 | 6.34 | 2.24 | 0.0774 |
| | day10 | male | f0 [Hz] | 6 | 206.76 | 193.86 | 43.98 | 17.95 | 0.117 |
| | | | Q | | 1.43 | 1.46 | 0.25 | 0.10 | 2e-3 (MWUS) |
| | | | Energy [kBT] | | 10.33 | 11.07 | 2.93 | 1.19 | 1e-3 (MWUS) |
| | day25 | male | f0 [Hz] | 19 | 274.37 | 274.50 | 48.26 | 11.07 | P = <0.001 (MWUS) |
| | | | Q | | 4.87 | 4.50 | 2.14 | 0.49 | P = <0.001 (MWUS) |
| | | | Energy [kBT] | | 33.31 | 29.47 | 13.70 | 3.14 | 0.012 (MWUS) |
| | | female | f0 [Hz] | 17 | 212.49 | 211.40 | 29.87 | 7.24 | 0.098 |
| | | | Q | | 2.79 | 2.29 | 1.28 | 0.31 | 0.121 |
| | | | Energy [kBT] | | 18.70 | 18.38 | 8.10 | 1.96 | 0.428 |
| | day50 | male | f0 [Hz] | 16 | 264.65 | 260.12 | 45.86 | 11.46 | 0.008 (MWUS) |
| | | | Q | | 2.32 | 1.99 | 1.44 | 0.36 | 0.234 |
| | | | Energy [kBT] | | 19.48 | 17.87 | 7.74 | 1.94 | 0.162 |
| | | female | f0 [Hz] | 20 | 220.15 | 230.52 | 35.89 | 8.02 | 0.0952 |
| | | | Q | | 1.83 | 1.69 | 0.67 | 0.15 | 0.124 |
| | | | Energy [kBT] | | 10.35 | 10.33 | 3.84 | 0.86 | P = <0.001 (MWUS) |
| | day60 | male | f0 [Hz] | 11 | 401.72 | 366.26 | 129.87 | 39.16 | P = <0.001 (MWUS) |
| | | | Q | | 1.44 | 1.28 | 0.38 | 0.12 | P = <0.001 (MWUS) |
| | | | Energy [kBT] | | 7.53 | 7.68 | 4.51 | 1.36 | P = <0.001 (MWUS) |
| | | female | f0 [Hz] | 12 | 243.38 | 234.37 | 29.08 | 8.39 | 4e-3 (MWUS) |
| | | | Q | | 1.25 | 1.14 | 0.38 | 0.11 | P = <0.001 (MWUS) |
| | | | Energy [kBT] | | 5.60 | 5.15 | 2.03 | 0.59 | P = <0.001 (MWUS) |
| | day70 | male | f0 [Hz] | 18 | 442.85 | 425.92 | 169.67 | 39.99 | P = <0.001 (MWUS) |

| | | | | | | | | | |
|---|---|---|---|---|---|---|---|---|---|
| | | | Q | | 1.09 | 1.04 | 0.32 | 0.08 | P = <0.001 (MWUS) |
| | | | Energy [kBT] | | 2.95 | 2.20 | 2.30 | 0.54 | P = <0.001 (MWUS) |
| | | female | f0 [Hz] | 17 | 439.19 | 359.96 | 195.84 | 47.50 | P = <0.001 (MWUS) |
| | | | Q | | 1.45 | 1.39 | 0.53 | 0.13 | 3e-3 (MWUS) |
| | | | Energy [kBT] | | 2.60 | 2.01 | 2.74 | 0.66 | P = <0.001 (MWUS) |
| OregonR | day1 | female | f0 [Hz] | 9 | 150.55 | 149.80 | 16.97 | 5.66 | -- |
| | | | Q | | 0.85 | 0.88 | 0.16 | 0.05 | -- |
| | | | Energy [kBT] | | 6.66 | 5.87 | 1.67 | 0.56 | -- |
| | day5 | female | f0 [Hz] | 8 | 147.30 | 139.40 | 29.16 | 10.31 | 0.78 |
| | | | Q | | 0.80 | 0.80 | 0.17 | 0.06 | 0.52 |
| | | | Energy [kBT] | | 5.13 | 4.46 | 2.03 | 0.72 | 0.061 |
| | day25 | female | f0 [Hz] | 5 | 164.01 | 160.66 | 69.37 | 31.02 | 0.79 |
| | | | Q | | 0.99 | 0.92 | 0.44 | 0.20 | 0.594 |
| | | | Energy [kBT] | | 7.14 | 5.30 | 3.58 | 1.60 | 0.732 |
| | day50 | female | f0 [Hz] | 10 | 481.62 | 544.13 | 201.75 | 63.80 | 4e-3 (MWUS) |
| | | | Q | | 0.70 | 0.72 | 0.19 | 0.06 | 0.0723 |
| | | | Energy [kBT] | | 1.14 | 0.76 | 1.19 | 0.38 | P = <0.001 (MWUS) |
| CantonS (G) | day1 | male | f0 [Hz] | 10 | 242.89 | 244.99 | 35.96 | 11.37 | -- |
| | | | Q | | 1.66 | 1.65 | 0.42 | 0.13 | -- |
| | | | Energy [kBT] | | 14.34 | 13.84 | 3.10 | 0.98 | -- |
| | | female | f0 [Hz] | 11 | 235.35 | 235.15 | 35.46 | 10.69 | -- |
| | | | Q | | 1.62 | 1.69 | 0.38 | 0.11 | -- |
| | | | Energy [kBT] | | 11.22 | 11.65 | 3.32 | 1.00 | -- |
| | day5 | male | f0 [Hz] | 12 | 268.96 | 273.22 | 28.19 | 8.14 | 0.071 |
| | | | Q | | 1.53 | 1.57 | 0.39 | 0.11 | 0.452 |
| | | | Energy [kBT] | | 9.27 | 9.34 | 2.46 | 0.71 | 3.64E-04 |
| | | female | f0 [Hz] | 9 | 264.02 | 253.02 | 42.16 | 14.05 | 0.116 |
| | | | Q | | 1.63 | 1.69 | 0.40 | 0.13 | 0.946 |
| | | | Energy [kBT] | | 7.41 | 7.88 | 3.24 | 1.08 | 0.0188 |
| | day25 | male | f0 [Hz] | 13 | 263.78 | 264.93 | 36.44 | 10.11 | 0.185 |
| | | | Q | | 1.69 | 1.52 | 0.52 | 0.14 | 0.905 |
| | | | Energy [kBT] | | 12.11 | 10.57 | 4.58 | 1.27 | 0.201 |
| | | female | f0 [Hz] | 12 | 253.57 | 256.22 | 32.53 | 9.39 | 0.213 |
| | | | Q | | 2.21 | 1.81 | 1.10 | 0.32 | 0.103 |
| | | | Energy [kBT] | | 10.73 | 9.15 | 4.39 | 1.27 | 0.765 |
| | day50 | male | f0 [Hz] | 12 | 370.57 | 342.20 | 88.07 | 25.42 | 3.63E-04 |
| | | | Q | | 1.37 | 1.22 | 0.47 | 0.14 | 0.142 |
| | | | Energy [kBT] | | 8.22 | 9.15 | 3.72 | 1.08 | 5.15E-04 |
| | | female | f0 [Hz] | 11 | 298.22 | 292.68 | 34.15 | 10.30 | 4.05E-04 |
| | | | Q | | 1.56 | 1.51 | 0.42 | 0.13 | 0.751 |
| | | | Energy [kBT] | | 8.28 | 6.40 | 3.38 | 1.02 | 0.0528 |
| | day68 | male | f0 [Hz] | 5 | 470.63 | 428.17 | 144.02 | 64.41 | 6e-3 (MWUS) |
| | | | Q | | 1.18 | 1.05 | 0.43 | 0.19 | 0.0608 |
| | | | Energy [kBT] | | 4.75 | 3.99 | 2.80 | 1.25 | 6.07E-05 |
| | | female | f0 [Hz] | 9 | 355.52 | 330.97 | 95.74 | 31.91 | P = <0.001 (MWUS) |

| | | | | | | | | | |
|---|---|---|---|---|---|---|---|---|---|
| | | | Q | | 2.17 | 2.06 | 0.95 | 0.32 | 0.224 |
| | | | Energy [kBT] | | 6.58 | 6.46 | 3.50 | 1.17 | 7.05E-03 |
| | day70 | male | f0 [Hz] | 8 | 590.29 | 588.41 | 167.25 | 59.13 | P = <0.001 (MWUS) |
| | | | Q | | 1.31 | 1.21 | 0.35 | 0.12 | 0.0797 |
| | | | Energy [kBT] | | 2.77 | 2.77 | 2.18 | 0.77 | 1.34E-07 |
| | | female | f0 [Hz] | 13 | 395.09 | 410.39 | 49.42 | 13.71 | P = <0.001 (MWUS) |
| | | | Q | | 1.60 | 1.69 | 0.47 | 0.13 | 0.922 |
| | | | Energy [kBT] | | 3.79 | 3.77 | 1.19 | 0.33 | P = <0.001 (MWUS) |
| CantonS (K) | day1 | male | f0 [Hz] | 12 | 256.60 | 255.85 | 26.09 | 7.53 | -- |
| | | | Q | | 1.53 | 1.64 | 0.27 | 0.08 | -- |
| | | | Energy [kBT] | | 10.59 | 11.51 | 2.88 | 0.83 | -- |
| | | female | f0 [Hz] | 10 | 213.18 | 213.91 | 34.64 | 10.95 | -- |
| | | | Q | | 1.41 | 1.33 | 0.46 | 0.14 | -- |
| | | | Energy [kBT] | | 9.05 | 9.45 | 2.51 | 0.79 | -- |
| | day5 | male | f0 [Hz] | 11 | 293.26 | 298.83 | 40.33 | 12.16 | 0.0163 |
| | | | Q | | 1.81 | 1.58 | 0.65 | 0.20 | 0.442 |
| | | | Energy [kBT] | | 8.03 | 7.62 | 2.36 | 0.71 | 0.0309 |
| | | female | f0 [Hz] | 10 | 252.48 | 234.85 | 56.75 | 17.94 | 0.054 |
| | | | Q | | 1.35 | 1.31 | 0.25 | 0.08 | 0.752 |
| | | | Energy [kBT] | | 5.88 | 5.61 | 1.82 | 0.57 | 4.63E-03 |
| | day25 | male | f0 [Hz] | 16 | 292.96 | 301.76 | 36.17 | 9.04 | 6.68E-03 |
| | | | Q | | 1.55 | 1.54 | 0.45 | 0.11 | 0.879 |
| | | | Energy [kBT] | | 6.48 | 6.77 | 2.58 | 0.65 | 5.16E-04 |
| | | female | f0 [Hz] | 11 | 288.89 | 288.09 | 10.98 | 3.31 | P = <0.001 (MWUS) |
| | | | Q | | 1.72 | 1.68 | 0.32 | 0.10 | 0.0785 |
| | | | Energy [kBT] | | 6.45 | 6.41 | 1.55 | 0.47 | 0.005 (MWUS) |
| | day50 | male | f0 [Hz] | 10 | 368.92 | 367.62 | 32.31 | 9.74 | 4.60E-09 |
| | | | Q | | 1.77 | 1.89 | 0.58 | 0.18 | 0.044 |
| | | | Energy [kBT] | | 3.88 | 3.99 | 1.41 | 0.43 | P = <0.001 (MWUS) |
| | | female | f0 [Hz] | 12 | 360.80 | 347.37 | 43.22 | 12.48 | 3.07E-08 |
| | | | Q | | 2.29 | 1.95 | 0.88 | 0.25 | 9.19E-03 |
| | | | Energy [kBT] | | 4.05 | 4.15 | 0.98 | 0.28 | 1e-3 (MWUS) |
| | day68 | male | f0 [Hz] | 7 | 377.56 | 388.55 | 35.42 | 13.39 | 1.44E-07 |
| | | | Q | | 1.95 | 2.11 | 0.46 | 0.17 | 0.0202 |
| | | | Energy [kBT] | | 3.92 | 3.99 | 1.12 | 0.42 | 2.04E-05 |
| | day60 | female | f0 [Hz] | 7 | 366.36 | 370.04 | 24.75 | 9.35 | 4.96E-08 |
| | | | Q | | 1.84 | 1.92 | 0.26 | 0.10 | 0.041 |
| | | | Energy [kBT] | | 3.60 | 3.35 | 1.06 | 0.40 | 4e-3 (MWUS) |
| | day70 | male | f0 [Hz] | 4 | 353.80 | 363.42 | 22.19 | 11.09 | 1.09E-05 |
| | | | Q | | 1.45 | 1.51 | 0.71 | 0.36 | 0.952 |
| | | | Energy [kBT] | | 3.81 | 4.33 | 2.04 | 1.02 | 7.16E-04 |

**Supplementary Table 5. 37 iRegulon predicted transcription factors (TFs) expressed in the *Drosophila* JO.** Genes that change their expression in one of the two age comparisons in both sexes are shown in bold type. The 19 genes functionally probed in our study are highlighted in light orange. 'Avg exp' stands for 'Average Expression'.

| Drosophila gene | Human orthologue | Avg exp | Description |
|---|---|---|---|
| Adf1 | no orthologs | 697.51 | Bess and MADF1 TF involved in the memory, learning and olfaction, and synapse formation |
| Aef1 | no orthologs | 560.58 | Zinc finger TF inhibits olfactory memory formation |
| **amos** | **ATOH1, ATOH7, NEUROD1** | 29.40 | bHlH TF involved in the sensory organ development. Amos specifies olfactory organ and the dendritic neurons in PNS |
| **aop** | **ETV6, 7** | 1032.95 | ETS and pnt TF involved to the lifespan determination. |
| **ara** | **IRX4, IRX6** | 315.33 | Iroquois complex TF controls patterning and proliferation |
| ato | ATOH1 | 1.66 | Developmental BHlH TF critical for chordotonal organs formation |
| Blimp-1 | PRDM1 | 1717.74 | Zinc finger and SET TF involved in the olfaction |
| ci | GLI1-3 | 512.22 | GLI C2H2-type zinc-finger TF belongs to Hh signalling pathway |
| ct | CUX1, CUX2 | 1099.71 | Developmental TF critical for JO formation |
| **Dr** | **MSX1,2** | 62.84 | TF involved in glial cell development suppressible by amos. Msx2 mutant mice show the optic nerve aplasia. |
| **Fd64A** | **FOXA1, A2, B1, B2 C1, C2, D2, D3, D4, D4L1, D4L3, D4L4, D4L5, D4L6, L2, E1, S1** | 25.06 | Fork-head DNA binding motif protein. FOXC1 is involved in the Axenfeld Rieger syndrome (sensorineural hearing loss) |
| ftz | HOXD4 | 12.42 | Homeobox domain TF controls segmentation |
| gl | ZNF500 | 405.85 | Zinc finger TF Characterized by Senthillian et al 2012 in sensory perception of sound |
| grh | GRHL1,2,3 | 1770.12 | Polycomb binding TF. GRHL2 involved in the autosomal dominant nonsyndromic deafness 28 |
| grn | GATA1-6 | 210.30 | Gata domain TF. GATA3 mutations showed sensorineural deafness in humans |
| Hr39 | no orthologs | 295.46 | nuclear hormone receptor and Zing finger/GATA like domain TF |
| Hsf | HSF1,2,4 | 657.92 | Heat-shock factor TF involved in defense response to the bacterial and fungal infection. HSF4 is involved in the cataract type 5 in humans. |

| | | | |
|---|---|---|---|
| **Jra** | **JUN, JUNB, JUND** | 655.83 | bZIP family, jun subfamily TF involved in JNK signalling and in the eye development |
| lola | ZBTB46, Bzel | 6102.76 | Zinc finger and BTB domain TF involved in axonogenesis, dendrite morphogenesis, locomotor activity and sound production |
| Med | SMAD4 | 327.42 | SMAD TF involved in BMP signalling pathway |
| nvy | CBFA2T3, 2, RUNX1T1 | 90.66 | Zinc finger TF enhances expression of Delta and Notch and involved in the morphogenesis of sensory neuron dendrites. |
| onecut | ONECUT1,2,3 | 875.91 | Cut domain TF expressed in a subset of olfactory neurons |
| Optix | SIX3, SIX6, SIX1 | 185.43 | SIX domain TF regulates dendrite morphogenesis and involved in eye formation. SIX1 is involved in the autosomal dominant deafness DFNA23. |
| p53 | P53, P63, P73 | 201.92 | Classic tumor-suppressor, involved in proliferation control. Suppressible by Dronc and CycE. |
| pho | YY2, ZFP42 | 883.01 | Zinc finger Plycomb TF involved in the regulation of many genes |
| **pnr** | **GATA1-6** | 103.37 | Gata domain TF |
| **Pph13** | **RAX, RAX2** | 92.26 | Homeobox TF expressed in all photoreceptors, upstream regulator of rhodopsins expression |
| **prd** | **PAX3, PAX5-8** | 95.58 | Paired and homeobox TF. Human PAX3 mutations showed CDHS (craniofacial-deafness-hand syndrome) |
| Rfx | RFX1,2,3 | 487.84 | Main regulator of mechanosensory ciliogenesis |
| run | RUNXA, RUNXB | 956.47 | TF required for axon guidance and dendrite morphodgenesis |
| **Sox100b** | **SOX8, SOX10** | 61.66 | Sox TF. Human SOX10 implicated in the Waardeburg syndrome (sensorineural hearing loss) |
| **Sox14** | **SOX12, SOX4, SOX11** | 900.65 | Sox TF involved in the gliogenesis in Drosophila, and the redox genes regulation |
| Sox21B . | SOX1, 2, 3, 4, 11, 12, 21 | 170.22 | Sox TF involved in olfactory behaviour, gliogenesis. |
| **srp** | **GATA4-6** | 250.13 | Gata domain TF involved into the haematopoiesis and immune system |
| Stat92E | STAT1, 3-6 | 1994.39 | Stat TF involved in Jak-STAT signalling pathway |
| **Tbp** | **TBP, TBPL2** | 188.21 | TATA binding TF. TBP involved in Parkinson disease. |

| | | | |
|---|---|---|---|
| **wor** | **SNAI2** | 56.76 | Zinc finger domain TF expressed in all neural tissues. SNAI2 involved in the Waardeburg syndrome type 2D |

**Supplementary Table 6. Auditory mechanics after overexpression (OE) or knockdown (KD) of transcriptional regulators.**

Three principal parameters of sound receiver function were assessed in gene KDs or OEs and their respective controls (shown in brackets): (i) f0 , the receivers' best frequency [in Hz]; (ii) the receivers' tuning sharpness or 'quality factor' Q [dimensionless] and (iii) the receivers' energy gain [in kBT]. Pair-wise student t-tests or Mann-Whitney (MWUS) tests were used to assess statistical significance (choice of test depending on data distribution). Significant changes are highlighted in yellow. Both males and females were measured, except for Optix OE, ato OE and Dhc98D KD, where only females were measured.

| Genotype | sex | Sample size | | Parameter | Mean | | Median | | StandardDev | | StandardErr | | p-value |
|---|---|---|---|---|---|---|---|---|---|---|---|---|---|
| Aef1 KD | male | 8 | (12) | f0 [Hz] | 253.04 | (247.24) | 251.87 | (236.40) | 28.65 | (42.66) | 10.13 | (12.86) | 0.743 |
| | | | | Q | 1.03 | (1.05) | 1.04 | (1.09) | 0.16 | (0.26) | 0.06 | (0.08) | 0.773 |
| | | | | Energy [kBT] | 5.71 | (7.27) | 5.48 | (7.27) | 1.64 | (1.76) | 0.58 | (0.53) | 0.0678 |
| | female | 10 | (11) | f0 [Hz] | 297.99 | (303.57) | 305.76 | (309.35) | 37.22 | (48.00) | 11.77 | (13.86) | 0.767 |
| | | | | Q | 1.30 | (1.41) | 1.28 | (1.38) | 0.27 | (0.34) | 0.09 | (0.10) | 0.433 |
| | | | | Energy [kBT] | 6.28 | (5.80) | 5.89 | (6.09) | 1.72 | (1.88) | 0.55 | (0.54) | 0.545 |
| amos KD | male | 9 | (7) | f0 [Hz] | 237.28 | (260.08) | 220.17 | (242.45) | 49.25 | (39.17) | 16.42 | (14.80) | 0.334 |
| | | | | Q | 0.85 | (1.18) | 0.87 | (1.09) | 0.16 | (0.30) | 0.05 | (0.11) | 6e-3 [MWRST] |
| | | | | Energy [kBT] | 3.97 | (7.19) | 4.62 | (6.56) | 1.54 | (2.06) | 0.51 | (0.78) | 3.05E-03 |
| | female | 9 | (8) | f0 [Hz] | 218.82 | (274.60) | 229.84 | (256.11) | 47.75 | (45.67) | 15.92 | (16.15) | 0.0269 |
| | | | | Q | 0.91 | (1.63) | 0.78 | (1.28) | 0.35 | (0.91) | 0.12 | (0.32) | 0.03 [MWUS] |
| | | | | Energy [kBT] | 1.95 | (4.01) | 2.03 | (4.06) | 0.55 | (0.99) | 0.18 | (0.35) | 7.38E-05 |
| aop KD | male | 6 | (11) | f0 [Hz] | 251.18 | (247.24) | 257.59 | (236.40) | 25.25 | (42.66) | 10.31 | (12.86) | 0.84 |
| | | | | Q | 0.90 | (1.05) | 0.83 | (1.09) | 0.28 | (0.26) | 0.11 | (0.08) | 0.276 |
| | | | | Energy [kBT] | 9.47 | (7.27) | 9.68 | (7.27) | 2.00 | (1.76) | 0.82 | (0.53) | 0.0326 |
| | female | 6 | (11) | f0 [Hz] | 279.39 | (309.11) | 280.63 | (310.65) | 32.80 | (46.14) | 13.39 | (13.91) | 0.185 |
| | | | | Q | 1.81 | (1.47) | 1.11 | (1.38) | 1.56 | (0.30) | 0.64 | (0.09) | 0.393 |
| | | | | Energy [kBT] | 7.09 | (5.67) | 7.09 | (5.66) | 3.02 | (1.92) | 1.23 | (0.58) | 0.252 |
| ara KD | male | 8 | (10) | f0 [Hz] | 292.99 | (266.39) | 293.97 | (265.83) | 28.72 | (47.35) | 10.15 | (14.97) | 0.083 |
| | | | | Q | 1.67 | (1.04) | 1.62 | (1.00) | 0.23 | (0.32) | 0.08 | (0.10) | 2.53E-04 |
| | | | | Energy [kBT] | 12.05 | (4.42) | 12.27 | (4.37) | 3.83 | (1.78) | 1.35 | (0.56) | 3.88E-05 |
| | female | 6 | (11) | f0 [Hz] | 259.80 | (282.12) | 267.75 | (275.95) | 24.28 | (32.67) | 9.91 | (9.85) | 0.165 |
| | | | | Q | 1.78 | (1.41) | 1.62 | (1.38) | 0.67 | (0.30) | 0.27 | (0.09) | 0.133 |
| | | | | Energy [kBT] | 9.52 | (4.48) | 9.46 | (3.93) | 2.94 | (1.53) | 1.20 | (0.46) | 3e-3 [MWRST] |
| ato KD | male | 12 | (11) | f0 [Hz] | 285.15 | (247.24) | 280.16 | (236.40) | 59.38 | (42.66) | 17.14 | (12.86) | 0.0958 |
| | | | | Q | 1.17 | (1.05) | 1.01 | (1.09) | 0.46 | (0.26) | 0.13 | (0.08) | 0.829 |
| | | | | Energy [kBT] | 6.38 | (7.27) | 5.63 | (7.27) | 2.71 | (1.76) | 0.78 | (0.53) | 0.364 |
| | female | 10 | (12) | f0 [Hz] | 273.07 | (303.57) | 279.71 | (309.35) | 36.26 | (48.00) | 11.47 | (13.86) | 0.114 |
| | | | | Q | 1.24 | (1.41) | 1.21 | (1.38) | 0.24 | (0.34) | 0.08 | (0.10) | 0.194 |
| | | | | Energy [kBT] | 4.50 | (5.80) | 4.39 | (6.09) | 1.06 | (1.88) | 0.34 | (0.54) | 0.0669 |
| ct KD | male | 8 | (9) | f0 [Hz] | 297.48 | (235.00) | 266.82 | (223.77) | 125.48 | (28.32) | 44.36 | (9.44) | 0.163 |
| | | | | Q | 1.21 | (0.99) | 1.06 | (0.97) | 0.38 | (0.25) | 0.13 | (0.08) | 0.16 |
| | | | | Energy [kBT] | 4.78 | (7.63) | 5.02 | (7.66) | 2.13 | (1.75) | 0.75 | (0.58) | 8.38E-03 |
| | female | 8 | (12) | f0 [Hz] | 290.15 | (303.57) | 296.88 | (309.35) | 39.29 | (48.00) | 13.89 | (13.86) | 0.52 |
| | | | | Q | 1.23 | (1.41) | 1.23 | (1.38) | 0.13 | (0.34) | 0.05 | (0.10) | 0.17 |
| | | | | Energy [kBT] | 4.76 | (5.80) | 4.42 | (6.09) | 1.90 | (1.88) | 0.67 | (0.54) | 0.245 |
| Dhc98D KD | female | 10 | (18) | f0 [Hz] | 347.14 | (276.94) | 282.40 | (284.77) | 183.24 | (59.16) | 57.94 | (13.94) | 0.719 |
| | | | | Q | 0.73 | (1.25) | 0.73 | (1.33) | 0.23 | (0.40) | 0.07 | (0.09) | 9.85E-04 |
| | | | | Energy [kBT] | 1.85 | (5.19) | 1.08 | (4.75) | 1.91 | (1.87) | 0.60 | (0.44) | 1.26E-04 |
| gl KD | male | 7 | [7) | f0 [Hz] | 310.56 | (257.56) | 326.21 | (263.27) | 34.71 | (45.77) | 13.12 | (16.18) | 0.0469 |
| | | | | Q | 1.03 | (0.91) | 1.06 | (0.96) | 0.18 | (0.19) | 0.07 | (0.07) | 0.164 |
| | | | | Energy [kBT] | 4.35 | (3.93) | 4.21 | (3.81) | 1.28 | (1.40) | 0.48 | (0.50) | 0.619 |

| | | | | | | | | | | | | | |
|---|---|---|---|---|---|---|---|---|---|---|---|---|---|
| | female | 5 | [7) | f0 [Hz] | 278.87 | (283.93) | 286.62 | (270.60) | 20.59 | (41.26) | 9.21 | (15.60) | 0.807 |
| | | | | Q | 1.07 | (1.40) | 1.06 | (1.32) | 0.20 | (0.35) | 0.09 | (0.13) | 0.0887 |
| | | | | Energy [kBT] | 2.20 | (4.50) | 2.27 | (3.88) | 0.83 | (1.61) | 0.37 | (0.61) | 0.0157 |
| lola KD | male | 5 | [9) | f0 [Hz] | 323.02 | (235.00) | 332.34 | (223.77) | 29.60 | (28.32) | 13.24 | (9.44) | 1.39E-04 |
| | | | | Q | 1.00 | (0.99) | 1.09 | (0.97) | 0.21 | (0.25) | 0.09 | (0.08) | 0.924 |
| | | | | Energy [kBT] | 5.33 | (7.63) | 5.25 | (7.66) | 1.80 | (1.75) | 0.80 | (0.58) | 0.0377 |
| | female | 6 | [11) | f0 [Hz] | 324.22 | (309.11) | 322.01 | (310.65) | 24.36 | (46.14) | 9.94 | (13.91) | 0.471 |
| | | | | Q | 1.14 | (1.47) | 1.14 | (1.38) | 0.09 | (0.30) | 0.04 | (0.09) | 0.0217 |
| | | | | Energy [kBT] | 5.09 | (5.67) | 5.36 | (5.66) | 1.18 | (1.92) | 0.48 | (0.58) | 0.58 |
| onecut KD | male | 10 | [11) | f0 [Hz] | 499.10 | (247.24) | 440.20 | (236.40) | 146.93 | (42.66) | 46.46 | (12.86) | P = <0.001 [MWRST] |
| | | | | Q | 1.07 | (1.05) | 0.96 | (1.09) | 0.32 | (0.26) | 0.10 | (0.08) | 0.846 |
| | | | | Energy [kBT] | 2.63 | (7.27) | 2.35 | (7.27) | 1.51 | (1.76) | 0.48 | (0.53) | 3.55E-06 |
| | female | 15 | [12) | f0 [Hz] | 420.61 | (303.57) | 407.05 | (309.35) | 37.42 | (48.00) | 9.66 | (13.86) | 1.81E-07 |
| | | | | Q | 0.88 | (1.41) | 0.90 | (1.38) | 0.16 | (0.34) | 0.04 | (0.10) | 1.55E-05 |
| | | | | Energy [kBT] | 1.50 | (5.80) | 1.58 | (6.09) | 0.54 | (1.88) | 0.14 | (0.54) | P = <0.001 [MWRST] |
| Optix KD | male | 10 | [11) | f0 [Hz] | 269.38 | (247.24) | 251.15 | (236.40) | 48.29 | (42.66) | 15.27 | (12.86) | 0.193 |
| | | | | Q | 1.45 | (1.05) | 1.34 | (1.09) | 0.33 | (0.26) | 0.10 | (0.08) | 5.14E-03 |
| | | | | Energy [kBT] | 7.30 | (7.27) | 7.39 | (7.27) | 1.94 | (1.76) | 0.61 | (0.53) | 0.971 |
| | female | 16 | [12) | f0 [Hz] | 243.94 | (303.57) | 241.04 | (309.35) | 25.29 | (48.00) | 6.32 | (13.86) | 0.000237 |
| | | | | Q | 1.55 | (1.41) | 1.48 | (1.38) | 0.51 | (0.34) | 0.13 | (0.10) | 0.562 |
| | | | | Energy [kBT] | 8.06 | (5.80) | 7.47 | (6.09) | 2.30 | (1.88) | 0.58 | (0.54) | 0.022 [MWUS] |
| pnr KD | male | 5 | [11) | f0 [Hz] | 314.09 | (247.24) | 319.14 | (236.40) | 37.71 | (42.66) | 16.87 | (12.86) | 9.54E-03 |
| | | | | Q | 1.28 | (1.05) | 1.21 | (1.09) | 0.30 | (0.26) | 0.13 | (0.08) | 0.137 |
| | | | | Energy [kBT] | 7.27 | (7.27) | 7.86 | (7.27) | 1.73 | (1.76) | 0.77 | (0.53) | 1 |
| | female | 5 | [12) | f0 [Hz] | 282.33 | (303.57) | 291.46 | (309.35) | 16.19 | (48.00) | 7.24 | (13.86) | 0.357 |
| | | | | Q | 1.37 | (1.41) | 1.48 | (1.38) | 0.41 | (0.34) | 0.18 | (0.10) | 0.822 |
| | | | | Energy [kBT] | 6.56 | (5.80) | 6.59 | (6.09) | 2.05 | (1.88) | 0.92 | (0.54) | 0.468 |
| Pph13 KD | male | 10 | [8) | f0 [Hz] | 278.64 | (257.56) | 276.56 | (263.27) | 36.82 | (45.77) | 11.64 | (16.18) | 0.294 |
| | | | | Q | 1.46 | (0.91) | 1.49 | (0.96) | 0.41 | (0.19) | 0.13 | (0.07) | 3.08E-03 |
| | | | | Energy [kBT] | 6.28 | (3.93) | 6.33 | (3.81) | 2.46 | (1.40) | 0.78 | (0.50) | 0.0289 |
| | female | 8 | [9) | f0 [Hz] | 297.15 | (282.57) | 298.02 | (270.60) | 45.70 | (36.45) | 16.16 | (12.15) | 0.476 |
| | | | | Q | 1.97 | (1.43) | 1.46 | (1.38) | 1.43 | (0.32) | 0.51 | (0.11) | 0.665 |
| | | | | Energy [kBT] | 6.64 | (4.52) | 6.11 | (3.88) | 2.54 | (1.70) | 0.90 | (0.57) | 0.0582 |
| Rfx KD | male | 6 | [11) | f0 [Hz] | 249.11 | (247.24) | 249.99 | (236.40) | 37.67 | (42.66) | 15.38 | (12.86) | 0.93 |
| | | | | Q | 1.09 | (1.05) | 0.94 | (1.09) | 0.31 | (0.26) | 0.13 | (0.08) | 0.761 |
| | | | | Energy [kBT] | 5.93 | (7.27) | 5.57 | (7.27) | 1.29 | (1.76) | 0.53 | (0.53) | 0.125 |
| | female | 7 | [12) | f0 [Hz] | 289.88 | (303.57) | 283.49 | (309.35) | 41.19 | (48.00) | 15.57 | (13.86) | 0.537 |
| | | | | Q | 1.43 | (1.41) | 1.34 | (1.38) | 0.43 | (0.34) | 0.16 | (0.10) | 0.923 |
| | | | | Energy [kBT] | 6.01 | (5.80) | 5.45 | (6.09) | 2.69 | (1.88) | 1.02 | (0.54) | 0.842 |
| run KD | male | 9 | [9) | f0 [Hz] | 303.89 | (235.00) | 290.25 | (223.77) | 40.60 | (28.32) | 13.53 | (9.44) | 2.00E-03 |
| | | | | Q | 1.52 | (0.99) | 1.58 | (0.97) | 0.49 | (0.25) | 0.16 | (0.08) | 8.00E-03 |
| | | | | Energy [kBT] | 9.10 | (7.63) | 8.76 | (7.66) | 4.29 | (1.75) | 1.43 | (0.58) | 0.377 |
| | female | 4 | [12) | f0 [Hz] | 286.85 | (303.57) | 295.09 | (309.35) | 40.44 | (48.00) | 20.22 | (13.86) | 0.543 |
| | | | | Q | 1.59 | (1.41) | 1.53 | (1.38) | 0.54 | (0.34) | 0.27 | (0.10) | 0.454 |

| | | | | | | | | | | | | | |
|---|---|---|---|---|---|---|---|---|---|---|---|---|---|
| | | | | Energy [kBT] | 6.63 | (5.80) | 5.81 | (6.09) | 3.08 | (1.88) | 1.54 | (0.54) | 0.52 |
| Sox14 KD | male | 5 | [11] | f0 [Hz] | 273.29 | (247.24) | 271.38 | (236.40) | 11.11 | (42.66) | 4.97 | (12.86) | 0.113 |
| | | | | Q | 1.22 | (1.05) | 1.26 | (1.09) | 0.13 | (0.26) | 0.06 | (0.08) | 0.187 |
| | | | | Energy [kBT] | 4.69 | (7.27) | 4.94 | (7.27) | 1.23 | (1.76) | 0.55 | (0.53) | 0.0109 |
| | female | 7 | [12] | f0 [Hz] | 277.09 | (303.57) | 268.76 | (309.35) | 55.07 | (48.00) | 20.82 | (13.86) | 0.287 |
| | | | | Q | 1.36 | (1.41) | 1.18 | (1.38) | 0.28 | (0.34) | 0.11 | (0.10) | 0.744 |
| | | | | Energy [kBT] | 4.55 | (5.80) | 4.92 | (6.09) | 1.64 | (1.88) | 0.62 | (0.54) | 0.164 |
| srp KD | male | 8 | [11] | f0 [Hz] | 284.02 | (247.24) | 287.25 | (236.40) | 31.45 | (42.66) | 11.12 | (12.86) | 0.0551 |
| | | | | Q | 1.12 | (1.05) | 0.99 | (1.09) | 0.34 | (0.26) | 0.12 | (0.08) | 0.634 |
| | | | | Energy [kBT] | 6.32 | (7.27) | 6.37 | (7.27) | 1.86 | (1.76) | 0.66 | (0.53) | 0.271 |
| | female | 6 | [12] | f0 [Hz] | 293.36 | (303.57) | 296.23 | (309.35) | 14.49 | (48.00) | 5.92 | (13.86) | 0.622 |
| | | | | Q | 1.23 | (1.41) | 1.18 | (1.38) | 0.28 | (0.34) | 0.11 | (0.10) | 0.282 |
| | | | | Energy [kBT] | 5.91 | (5.80) | 5.35 | (6.09) | 1.00 | (1.88) | 0.41 | (0.54) | 0.963 |
| Stat92E KD | male | 9 | [11] | f0 [Hz] | 286.36 | (247.24) | 300.36 | (236.40) | 48.83 | (42.66) | 16.28 | (12.86) | 0.0718 |
| | | | | Q | 0.98 | (1.05) | 0.91 | (1.09) | 0.19 | (0.26) | 0.06 | (0.08) | 0.504 |
| | | | | Energy [kBT] | 5.87 | (7.27) | 6.08 | (7.27) | 2.18 | (1.76) | 0.73 | (0.53) | 0.128 |
| | female | 7 | [12] | f0 [Hz] | 302.75 | (303.57) | 301.52 | (309.35) | 20.59 | (48.00) | 7.78 | (13.86) | 0.967 |
| | | | | Q | 1.11 | (1.41) | 1.11 | (1.38) | 0.18 | (0.34) | 0.07 | (0.10) | 0.0484 |
| | | | | Energy [kBT] | 4.96 | (5.80) | 4.82 | (6.09) | 0.82 | (1.88) | 0.31 | (0.54) | 0.281 |
| Tbp KD | male | 10 | [7] | f0 [Hz] | 254.29 | (260.08) | 234.91 | (242.45) | 58.22 | (39.17) | 18.41 | (14.80) | 0.823 |
| | | | | Q | 1.19 | (1.18) | 1.09 | (1.09) | 0.35 | (0.30) | 0.11 | (0.11) | 0.98 |
| | | | | Energy [kBT] | 5.82 | (7.19) | 5.47 | (6.56) | 2.55 | (2.06) | 0.81 | (0.78) | 0.256 |
| | female | 9 | [8] | f0 [Hz] | 246.73 | (274.60) | 248.12 | (256.11) | 21.39 | (45.67) | 7.13 | (16.15) | 0.121 |
| | | | | Q | 1.31 | (1.63) | 1.23 | (1.28) | 0.44 | (0.91) | 0.15 | (0.32) | 0.532 |
| | | | | Energy [kBT] | 4.09 | (4.01) | 3.75 | (4.06) | 1.70 | (0.99) | 0.57 | (0.35) | 0.905 |
| wor KD | male | 7 | [8] | f0 [Hz] | 297.67 | (257.56) | 300.41 | (263.27) | 36.91 | (45.77) | 13.95 | (16.18) | 0.0873 |
| | | | | Q | 1.77 | (0.91) | 1.66 | (0.96) | 0.35 | (0.19) | 0.13 | (0.07) | P= <0.001 [MWRST] |
| | | | | Energy [kBT] | 12.36 | (3.93) | 10.74 | (3.81) | 4.40 | (1.40) | 1.66 | (0.50) | P = <0.001 [MWRST] |
| | female | 14 | [15] | f0 [Hz] | 266.28 | (277.04) | 259.42 | (275.95) | 37.25 | (30.82) | 9.95 | (7.96) | 0.403 |
| | | | | Q | 2.59 | (1.41) | 1.99 | (1.38) | 1.65 | (0.31) | 0.44 | (0.08) | P = <0.001 [MWRST] |
| | | | | Energy [kBT] | 9.75 | (4.31) | 8.66 | (3.88) | 3.87 | (1.48) | 1.04 | (0.38) | P = <0.001 [MWRST] |
| amos OE [30d] | male | 4 | [6] | f0 [Hz] | 279.72 | (419.64) | 286.13 | (399.99) | 71.42 | (98.17) | 35.71 | (40.08) | 0.041 |
| | | | | Q | 1.08 | (0.88) | 1.01 | (0.84) | 0.35 | (0.29) | 0.17 | (0.12) | 0.365 |
| | | | | Energy [kBT] | 8.80 | (1.20) | 8.95 | (1.05) | 3.60 | (0.59) | 1.80 | (0.24) | P = 0.01 [MWRST] |
| | female | 9 | [10] | f0 [Hz] | 304.84 | (336.28) | 298.36 | (326.49) | 46.85 | (45.84) | 15.62 | (14.50) | 0.158 |
| | | | | Q | 1.25 | (0.93) | 1.18 | (0.93) | 0.34 | (0.22) | 0.11 | (0.07) | 0.0253 |
| | | | | Energy [kBT] | 7.03 | (1.13) | 6.91 | (1.11) | 2.55 | (0.71) | 0.85 | (0.22) | P = <0.001 [MWRST] |
| Optix OE [15d] | male | 4 | [3] | f0 [Hz] | 591.51 | (320.86) | 606.59 | (313.55) | 115.18 | (21.63) | 57.59 | (12.49) | 0.0111 |
| | | | | Q | 0.94 | (1.23) | 0.91 | (1.26) | 0.20 | (0.55) | 0.10 | (0.32) | 0.629 |
| | | | | Energy [kBT] | 0.43 | (5.88) | 0.44 | (6.85) | 0.25 | (3.10) | 0.12 | (1.79) | 0.0152 |
| | female | 5 | [8] | f0 [Hz] | 355.45 | (278.15) | 386.37 | (280.29) | 62.44 | (21.54) | 27.92 | (7.62) | 7.38E-03 |
| | | | | Q | 1.21 | (1.76) | 1.28 | (1.75) | 0.39 | (0.40) | 0.18 | (0.14) | 0.0346 |

| | | | | | | | | | | | | | |
|---|---|---|---|---|---|---|---|---|---|---|---|---|---|
| | | | | Energy [kBT] | 1.87 | (6.94) | 2.26 | (6.50) | 1.12 | (3.16) | 0.50 | (1.12) | 5.83E-03 |
| Optix OE [30d] | female | 6 | [10) | f0 [Hz] | 449.41 | (336.28) | 465.23 | (326.49) | 36.54 | (45.84) | 14.92 | (14.50) | 1.55E-04 |
| | | | | Q | 1.27 | (0.93) | 1.24 | (0.93) | 0.24 | (0.22) | 0.10 | (0.07) | 0.0125 |
| | | | | Energy [kBT] | 1.22 | (1.13) | 1.20 | (1.11) | 0.65 | (0.71) | 0.26 | (0.22) | 0.807 |
| wor OE [15d] | male | 8 | [10) | f0 [Hz] | 304.60 | (266.39) | 296.64 | (265.83) | 26.10 | (47.35) | 9.23 | (14.97) | 0.0582 |
| | | | | Q | 1.27 | (1.04) | 1.23 | (1.00) | 0.24 | (0.32) | 0.09 | (0.10) | 0.116 |
| | | | | Energy [kBT] | 5.77 | (4.42) | 5.00 | (4.37) | 1.87 | (1.78) | 0.66 | (0.56) | 0.139 |
| | female | 6 | [15) | f0 [Hz] | 261.36 | (277.04) | 236.75 | (275.95) | 60.68 | (30.82) | 24.77 | (7.96) | 0.437 |
| | | | | Q | 1.19 | (1.41) | 1.07 | (1.38) | 0.45 | (0.31) | 0.18 | (0.08) | 0.208 |
| | | | | Energy [kBT] | 4.46 | (4.31) | 4.00 | (3.88) | 1.15 | (1.48) | 0.47 | (0.38) | 0.669 |
| gl OE [30d] | male | 4 | [6) | f0 [Hz] | 388.92 | (419.64) | 404.10 | (399.99) | 58.65 | (98.17) | 29.33 | (40.08) | 0.593 |
| | | | | Q | 0.69 | (0.88) | 0.71 | (0.84) | 0.15 | (0.29) | 0.07 | (0.12) | 0.255 |
| | | | | Energy [kBT] | 0.96 | (1.20) | 0.99 | (1.05) | 0.26 | (0.59) | 0.13 | (0.24) | 0.458 |
| | female | 4 | [7) | f0 [Hz] | 214.44 | (326.72) | 211.58 | (321.16) | 16.30 | (49.25) | 8.15 | (18.61) | 1.88E-03 |
| | | | | Q | 0.51 | (0.95) | 0.48 | (0.96) | 0.10 | (0.19) | 0.05 | (0.07) | 6.00E-03 |
| | | | | Energy [kBT] | 0.95 | (1.36) | 1.07 | (1.49) | 0.28 | (0.74) | 0.14 | (0.28) | 0.323 |
| ato OE | female | 9 | [10) | f0 [Hz] | 299.31 | (336.28) | 290.41 | (326.49) | 115.45 | (45.84) | 38.48 | (14.50) | 0.54 |
| | | | | Q | 1.34 | (0.93) | 0.99 | (0.93) | 0.81 | (0.22) | 0.27 | (0.07) | 0.348 |
| | | | | Energy [kBT] | 4.36 | (1.13) | 4.56 | (1.11) | 1.71 | (0.71) | 0.57 | (0.22) | P = <0.001 [MWRST] |
